# Roles of Kdm6a and Kdm6b in regulation of mammalian neural regeneration

**DOI:** 10.1101/2023.09.12.557354

**Authors:** Shu-Guang Yang, Xue-Wei Wang, Chang-Ping Li, Tao Huang, Cheng Qian, Qiao Li, Ling-Rui Zhao, Si-Yu Zhou, Chen-Yun Ding, Saijilafu, Chang-Mei Liu, Feng-Quan Zhou

**Affiliations:** Center for translational neural regeneration research, Sir Run Run Shaw Hospital, Zhejiang University School of Medicine, Hangzhou, Zhejiang 310016, China; Department of Orthopedic Surgery, The Johns Hopkins University School of Medicine, Baltimore, MD 21205, USA; The Solomon H. Department of Neuroscience, The Johns Hopkins University School of Medicine, Baltimore, MD 21205, USA; Department of Molecular Medicine, University of South Florida Morsani College of Medicine, Tampa, Florida, USA; Key Laboratory of Organ Regeneration and Reconstruction, Institute of Zoology, Chinese Academy of Sciences; Institute for Stem Cell and Regeneration, Chinese Academy of Sciences; Beijing Institute for Stem Cell and Regenerative Medicine, Beijing 100101, China; Savaid Medical School, University of Chinese Academy of Sciences, Beijing 100049, China; State Key Laboratory of Frigid Zone Cardiovascular Diseases (SKLFZCD), Department of Cardiovascular Surgery, General Hospital of Northern Theater Command, Shenyang, Liaoning 110016, China; Key Laboratory of Novel Targets and Drug Study for Neural Repair of Zhejiang Province, School of Medicine, Hangzhou City University, Hangzhou, Zhejiang 310015, China

**Keywords:** Kdm6a, Kdm6b, H3K27me3, optic nerve regeneration, neuroprotection, sensory axon regeneration, epigenetic regulation

## Abstract

Epigenetic regulation of neuronal transcriptomic landscape is emerging to be a key coordinator of mammalian neural regeneration. Here we investigated roles of two histone 3 lysine 27 (H3K27) demethylases Kdm6a/b in controlling neuroprotection and axon regeneration. Deleting either Kdm6a or Kdm6b led to enhanced sensory axon regeneration in the PNS, whereas in the CNS only deleting Kdm6a in retinal ganglion cells (RGCs) significantly enhanced optic nerve regeneration. Moreover, both Kdm6a and Kdm6b functioned to regulate RGC survival but with different mechanisms. Mechanistically, Kdm6a regulates RGC regeneration via distinct pathway from that of Pten and co-deleting Kdm6a and Pten resulted in long distance optic nerve regeneration passing the optic chiasm. In addition, RNA-seq profiling revealed that Kdm6a deletion switched the RGC transcriptomics into a developmental-like state and suppressed several known repressors of neural regeneration. Klf4 was identified as a direct downstream target of Kdm6a-H3K27me3 signaling in both sensory neurons and RGCs to regulate axon regeneration. These findings not only revealed different roles of Kdm6a and Kdm6b in regulation of neural regeneration and their underlying mechanisms, but also identified Kdm6a- mediated histone demethylation signaling as a novel epigenetic target for supporting CNS neural regeneration.

## 1. Introduction

Axon regeneration in the mammalian central nervous system (CNS) has been a long standing and highly challenging issue in the biomedical research field. A major reason for the failed CNS axon regeneration is that mature mammalian CNS neurons permanently lose their intrinsic ability to support axon growth during maturation.[1–3] Recent studies have identified many genes that could be manipulated to significantly boost the intrinsic axon regeneration ability of mature CNS neurons, including *Pten*, *Klf4/6/7/9*, *Gsk3b*, *Socs3*, *c-Myc*, *B-Raf, and Lin28.*[4–12] However, the molecular mechanisms, especially at the transcriptional and chromatin regulation level, by which many genes act coordinately to regulate axon regeneration is still rudimentary and fragmented. Thus, understanding the intrinsic transcriptomic networks and chromatin modulation specifically underlying axon growth during development and regeneration is of great importance in the neural regeneration field.

Epigenetic regulation is a key mechanism to control chromatin structure and the subsequent transcriptomic state,[13] making it a highly functional mechanism for coordinating gene transcription during axon regeneration. For instance, several studies, including ours,[11, 14–19] have illustrated the important roles of microRNAs in regulation of PNS axon regeneration. In addition, DNA methylation has also been shown to be involved in axon regeneration.[20–22] Dynamic histone acetylation could control chromatin accessibility and gene transcription of regeneration-associated genes (RAGs) to support the regenerative capacity.[23–26] Moreover, our study[16] has provided *in vivo* evidence that microRNA-138 and histone deacetylase Sirt1 form a mutual negative feedback loop to control sensory axon regeneration, suggesting crosstalk between two epigenetic signaling. Furthermore, a recent study[27] using multiomics sequencing characterized the unique chromatin signature and associated transcription profile during spontaneous sensory axon regeneration, highlighting the key roles of histone acetylation in supporting axon regeneration. Although these previous studies provided clear evidence about the involvement of epigenetic regulation in neural regeneration, to our knowledge, to date very few studies[28, 29] have demonstrated that manipulation of a single epigenetic regulator alone is able to induce long-distance CNS axon regeneration.

Tri-methylation of histone 3 lysine 27 (H3K27me3) functions to repress transcription of selected genes. Our recent studies investigated the roles of H3K27 methyltransferase Ezh2 in neural development and regeneration, revealing Ezh2 as a major factor shaping neural structure. Specifically, we showed that in postmitotic neurons Ezh2 acted to control multiple steps of neuronal morphogenesis during development, thereby leading to cognitive consequences in adult mice.[30] Our latest study[28] showed that Ezh2 acted to support long-distance axon regeneration in both PNS and CNS via coordinating gene transcription in multiple regenerative pathways. To date, three H2K27 demethylases have been identified, including Lysine (K)- specific demethylase 6a encoded by *Kdm6a* (or *Utx*) located on the X chromosome, Kdm6b encoded by *Kdm6b* (or *Jmjd3*), and Kdm6c encoded by *Kdm6c* (or *Uty*) located on the Y chromosome. They act specifically to erase trimethyl groups of H3K27me3, thereby generating transcriptionally permissive chromatin.[31, 32] As the paralog of Kdm6a, Kdm6c has been shown to have much reduced enzymatic activity.[31, 33, 34] The roles and mechanisms of both Kdm6a and Kdm6b have been extensively studied for their involvement in development,[31, 32, 35] differentiation,[36, 37] cell plasticity,[38] aging,[39–42] cell reprogramming,[43, 44] inflammatory response,[45, 46] tumorigenesis,[47, 48] and neurodegeneration.[49, 50] Previous studies have shown that Kdm6a and Kdm6b could play either similar or distinct roles in different biological processes. For instance, in mouse spinal motor neurons Kdm6b but not Kdm6a functions to regulate subtype diversification during development.[51] In contrast, Kdm6a but not Kdm6b is able to act as a direct sensor for cellular oxygen level and subsequent changes in chromatin and cell fate.[52] In the nervous system, our recent study[53] demonstrated that Kdm6a functioned to regulate neuronal dendritic development and various cognitive functions through demethylation of H3K27me3. In addition, previous studies suggested that Kdm6b served as a crucial regulator of neuronal maturation and survival.[54, 55] There is early evidence that PNS axotomy can trigger dynamic changes of histone methylation in sensory neurons.[26] However, to date, the functional role of histone methylation in axon regeneration is still under intensive investigation.

Here, we report that deleting *Kdm6a* or *Kdm6b* significantly promotes sensory axon regeneration *in vivo*. Importantly, deleting *Kdm6a*, but not *Kdm6b,* in retinal ganglion cells (RGCs) markedly enhanced optic nerve regeneration. Mechanistically, our data demonstrated that Kdm6a acted via a Pten/PI3K independent pathway to enhance optic nerve regeneration. Moreover, by analyzing and comparing transcriptomic changes of purified RGCs during development/maturation to that of *Kdm6a* knockout, we found that deleting *Kdm6a* reactivated developmental-like growth programs via modified chromatin structure and transcriptomics in RGCs. Moreover, we identified several known axon growth repressor genes as downstream targets suppressed by the Kdm6a-H3K27me3 signaling. In particular, we provided strong evidence that the transcription factor Klf4 was directly regulated by H3K27me3, and overexpressing Klf4 in adult sensory neurons significantly blocked axon regeneration *in vivo*. In addition, deletion of either Kdm6a or Kdm6b markedly increased RGC survival, and co-deleting Kdm6a/b exhibited additive effect on RGC protection, indicating their distinct underlying mechanisms. Collectively, our findings revealed that Kdm6a played key roles in repressing either PNS or CNS axon regeneration, whereas Kdm6b was not involved in regulating CNS axon regeneration. Moreover, both demethylases were able to regulate CNS neuronal survival after injury, but with non-overlapping mechanisms. Finally, we provided evidence that deleting Kdm6a functioned to enhance mammalian CNS neural regeneration by reshaping the neuronal transcriptomic landscape to a developmental-like state. Our study not only characterized the novel but non-overlapping roles of Kdm6a and Kdm6b in regulation of mammalian neural regeneration, but also discovered promising new molecular targets for promoting neuroprotection and axon regeneration.

## 2. Results

### 2.1. Kdm6a and Kdm6b are negative regulators of spontaneous sensory axon regeneration

To characterize the roles of demethylases Kdm6a/b in axon injury and regeneration, we first examined how they acted in sensory neurons in the dorsal root ganglion (DRG) during spontaneous axon regeneration. Time-course analyses by real-time PCR showed that the mRNA levels of *Kdm6a* were gradually downregulated after sciatic nerve injury (SNI), reaching its minimum at day 7 post-SNI (**Figure 1a**). Similarly, the levels of *Kdm6b* were also significantly reduced 1-day post-SNI but recovered back at day 3 and 7 (**Figure 1a**). We then performed immunostaining analyses to further examine the protein levels of Kdm6a or Kdm6b in sensory neurons *in vivo*. Confocal imaging of DRG sections showed that Kdm6a and Kdm6b were presented in both the nucleus and cytoplasm of sensory neurons before SNI. After SNI, the protein levels of Kdm6a in both the nuclei and cytoplasm of sensory neurons were significantly reduced 1 and 3 days later (**Figure 1b, c**). Kdm6b staining demonstrated different change patterns after SNI with its protein level only reduced in both the nucleus and cytoplasm 1 day after SNI. In day 3 post SNI, the level of Kdm6b was restored in the cytoplasm but kept low in the nucleus (**Figure 1d, e**). As histone demethylases, both proteins function in the nucleus. Thus, these data suggested that Kdm6a/b might act to suppress sensory axon regeneration. We therefore tested if downregulation of Kdm6a or Kdm6b in sensory neurons could enhance axon regeneration *in vivo*. We used our well-established *in vivo* electroporation technique[56] to co-transfect sensory neurons with a group of 4 different siRNAs against *Kdm6a* or *Kdm6b* and the EGFP plasmid. According to our previous study, the transfection efficiency of fluorescence labeled siRNAs and EGFP via *in vivo* electroporation is over 96%,[11] and 5-10%,[56] respectively. Therefore, we considered all EGFP positive neurons as siRNA positive. Western blot results showed that siRNAs significantly reduced the protein level of Kdm6a or Kdm6b *in vivo* (**Figure 1f, g**). To assess the functional roles of Kdm6a or Kdm6b in sensory axon regeneration, we performed SNI 2 days after electroporation and analyzed axon regeneration 2 days post-SNI when spontaneous sensory axon regeneration was just starting (**Figure 1h**). The results showed that knocking down *Kdm6a* or *Kdm6b* in sensory neurons markedly enhanced axon regeneration *in vivo* (**Figure 1i-l**). Together, these results demonstrated clearly that demethylases Kdm6a/b function similarly as important repressors of sensory axon regeneration.

**Fig. 1.**
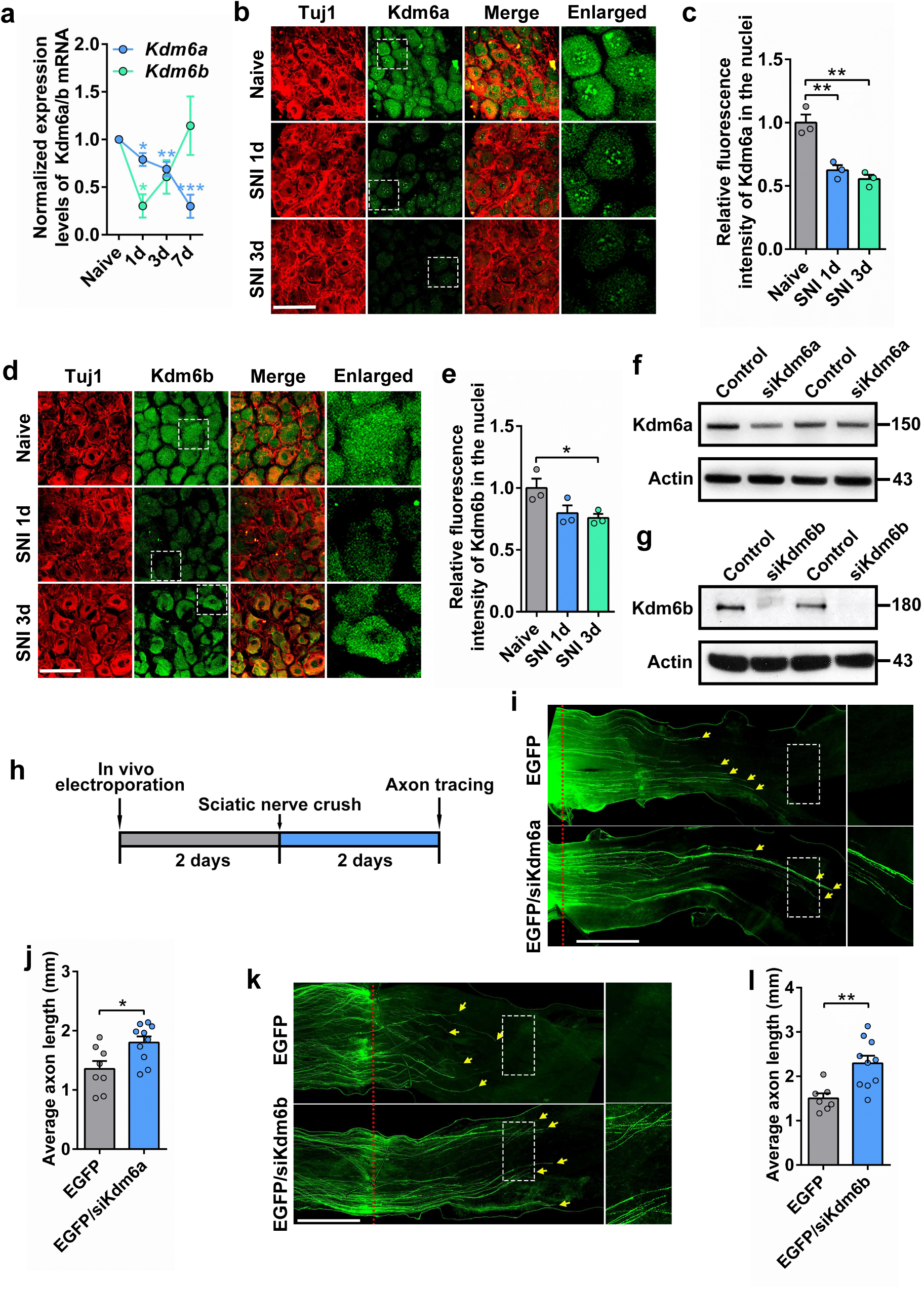
Knocking down *Kdm6a* or *Kdm6b* in adult mouse sensory neurons markedly enhanced sensory axon regeneration in vivo. (a) Time course of expression of *Kdm6a* and *Kdm6b* transcripts in ipsilateral L4/L5 DRG from naive mice and 1, 3, and 7 days post-sciatic nerve crush by real time PCR analysis. (one way ANOVA followed by Tukey’s multiple comparison test, *Kdm6a*: *P*=0.0001, n=5 independent experiments for each condition; *Kdm6b*: *P*=0.0165, n=7 independent experiments for each condition). (b) Immunostaining of DRG sections showing significantly reduced level of Kdm6a in adult mouse sensory neurons in vivo after sciatic nerve injury (SNI). The 4^th^ column shows enlarged images of areas indicated by the dashed white boxes in the 2^nd^ column. Images were stained for neuronal marker Tuj1 (red) and Kdm6a (green). Scale bar, 100 µm for the first 3 columns and 30 µm for the 4^th^ column. (c) Quantification of relative fluorescence intensity of Kdm6a immunostaining shown in (b) (two tailed student’s *t* tests, Naïve vs SNI 1d: *P*= 0.0076, Naïve vs SNI 3d: *P*= 0.0036; n= 3 independent experiments for each condition). (d) Immunostaining of DRG sections showing significantly reduced level of Kdm6b in adult mouse sensory neurons in vivo at day 3 after sciatic nerve injury (SNI). The 4^th^ column shows enlarged images of areas indicated by the dashed white boxes in the 2^nd^ column. Images were stained for neuronal marker Tuj1 (red) and Kdm6b (green). Scale bar, 100 µm for the first 3 columns and 30 µm for the 4^th^ column. (e) Quantification of relative fluorescence intensity of Kdm6b immunostaining shown in (d) (two tailed student’s *t* tests, Naïve vs SNI 3d: *P*=0.0426; n=3 independent experiments for each condition). (f) Western blot image showing greatly reduced protein level of Kdm6a in DRG tissues 2 days after in vivo electroporation of siRNAs against *Kdm6a* (siKdm6a). A representative image from 2 independent experiments is shown. (g) Western blot image showing greatly reduced protein level of Kdm6b in DRG tissues 2 days after in vivo electroporation of siRNAs against *Kdm6b* (siKdm6b). A representative image from 2 independent experiments is shown. (h) Schematics of in vivo electroporation and investigation of axon regeneration after knocking down *Kdm6a* or *Kdm6b*. L4 and L5 DRGs of wild type adult mice were electroporated in vivo with EGFP or EGFP + siRNAs against *Kdm6a* (siKdm6a) or *Kdm6b* (siKdm6b). To assess the promoting effects, axon regeneration analyses were performed 2 days post-sciatic nerve crush, when sensory axon regeneration has not reached its optimal rate. (i) Representative images of in vivo sensory axon regeneration in mice electroporated with either EGFP or EGFP+siKdm6a. The right column shows enlarged images of areas indicated by the dashed white boxes in the left column. The red dotted lines indicate the nerve crush sites. Yellow arrows indicate the distal ends of regenerating axons. Scale bar, 1 mm for the left column and 0.5 mm for the right column. (j) Knocking down *Kdm6a* in sensory neurons significantly promoted sensory axon regeneration in vivo (two tailed student’s *t* tests, *P*=0.0152; EGFP; n= 8 mice, EGFP+siKdm6a; n= 10 mice). (k) Representative images of in vivo sensory axon regeneration in mice electroporated with either EGFP or EGFP+siKdm6b. The right column shows enlarged images of areas indicated by the dashed white boxes in the left column. The red dotted lines indicate the nerve crush sites. Yellow arrows indicate the distal ends of regenerating axons. Scale bar, 1 mm for the left column and 0.5 mm for the right column. (l) Knocking down *Kdm6b* in sensory neurons significantly promoted sensory axon regeneration in vivo (two tailed student’s *t* tests, *P*=0.0033; EGFP: n=7 mice, EGFP+siKdm6b: n=10 mice). *, **, \*\*\**P* < 0.05, 0.01, 0.001, respectively. Data are represented as mean ± SEM.

### 2.2. *Kdm6a* knockout enhances optic nerve regeneration and RGC survival

Since the role of Kdm6a in sensory axon regeneration has been confirmed, we explored how Kdm6a regulated CNS neural regeneration using the optic nerve regeneration model. To examine the mRNA levels of *Kdm6a* in RGCs, we enriched the Thy1.2 positive RGCs by immunomagnetic cell separation (MACS) and the positive rate of Thy1.2 in positive fraction was 80.8±4.43% based on flow cytometric analysis (**Extended Data** Figure 1a**, b**). Unlike that of sensory neurons, real-time PCR analysis showed that the mRNA levels of *Kdm6a* in RGCs were not changed by optic nerve crush (ONC) (**Extended Data** Figure 1c). We next performed immunostaining analysis to examine the protein levels of Kdm6a in RGCs after ONC. The results showed that Kdm6a was highly enriched in the nuclei of RGCs and its level was unchanged after ONC (**Figure 2a-c**), confirming the real-time PCR results. By using an available single-cell RNA-seq (scRNA-seq) dataset of adult mouse RGCs,[57] we further confirmed that the expression levels of Kdm6a in RGCs were unchanged at different time points after ONC (**Extended Data** Figure 2a**, b**). The unchanged levels of Kdm6a in RGCs upon ONC are in stark contrast to that occurred in sensory neurons upon SNI, suggesting that Kdm6a in RGCs acts to suppress optic nerve regeneration.

**Fig. 2.**
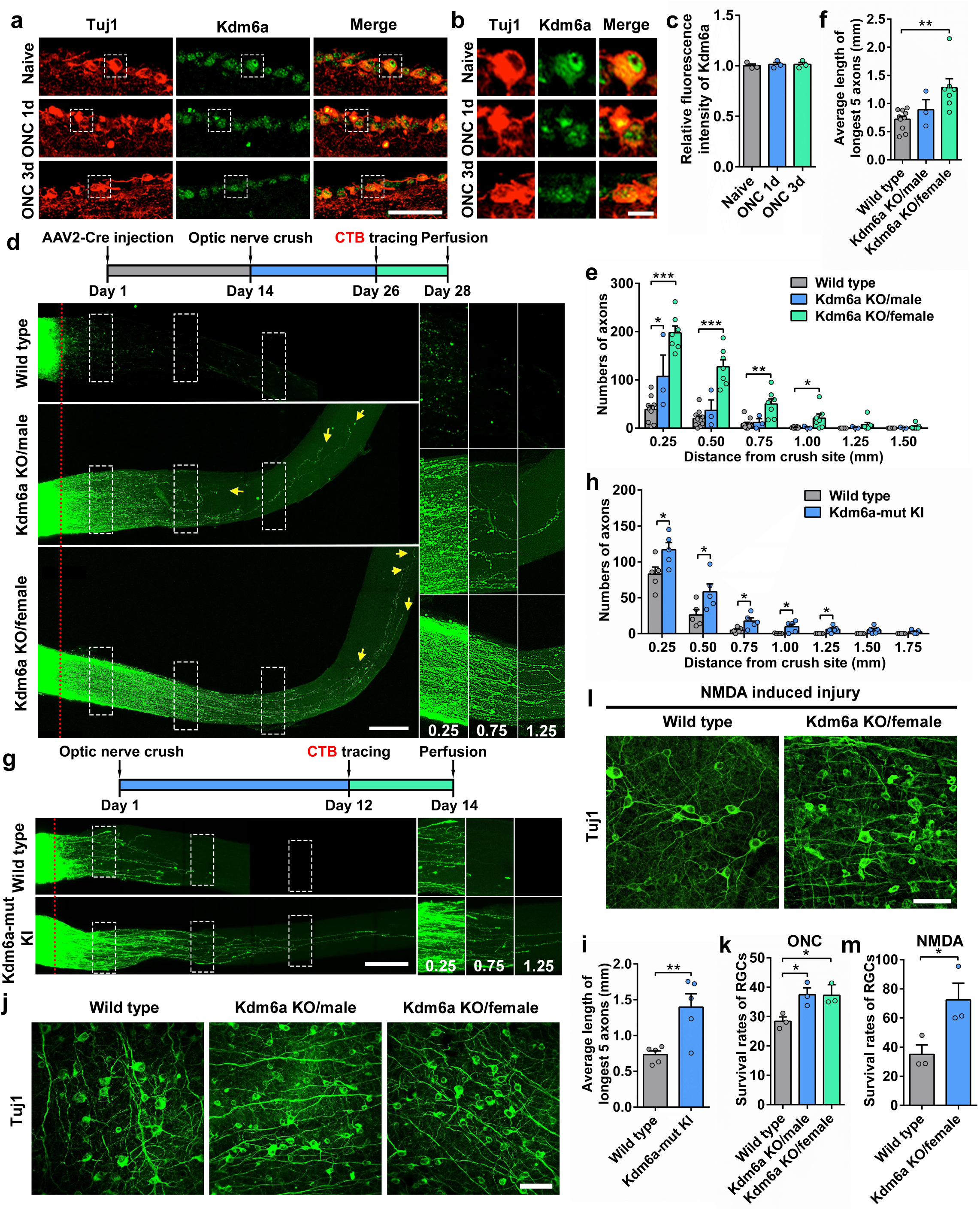
**Conditional deletion of H3K27 demethylase Kdm6a in mouse retinal ganglion cells (RGCs) markedly promoted optic nerve regeneration and RGC survival.** (a) Representative images of Kdm6a immunostaining in retina sections showing unchanged level of Kdm6a in RGCs after optic nerve crush (ONC). Images were stained for neuronal marker Tuj1 (red) and Kdm6a (green). Scare bar, 50 μm. (b) Enlarged images of areas indicated by the dashed white boxes in (a). Scale bar, 10 µm. (c) Quantification of relative fluorescence intensity of Kdm6a immunostaining in (a) (one way ANOVA followed by Tukey’s multiple comparison test, P=0.2274; n=3 independent experiments for each condition). (d) Top: schematic time course of optic nerve regeneration model. Specifically, AAV2-Cre was injected at day 1 and the optic nerve was crushed 2 weeks later. Optic nerve regeneration was analyzed 2 weeks after the nerve crush, and the fluorescence (Alexa 594) tagged cholera toxin β subunit (CTB) was injected 2 days before collecting the optic nerve. Bottom: representative 2D projected confocal images of whole mount cleared optic nerves of wild type, male *Kdm6a* knockout (Kdm6a KO/male), female *Kdm6a* knockout (Kdm6a KO/female) mice 2 weeks after ONC. The regenerating axons were labeled with CTB-Alexa 594. Yellow arrows indicate the distal ends of regenerating axons. The red dotted lines indicate the nerve crush sites. The columns on the right display magnified images of the areas in white, dashed boxes on the left, showing axons at 0.25, 0.75, and 1.25 mm distal to the crush sites. Scale bar, 200 µm for the left column and 100 µm for the magnified images. (e) Quantification of regenerating axons at different distances distal to the nerve crush site (0.25 to 1.50 mm) (one way ANOVA followed by Tukey’s multiple comparisons test, *P<*0.0001; wild type: n=10 mice, Kdm6a KO/ male: n=3 mice, Kdm6a KO/female: n=7). (f) Quantification of the average length of the top 5 longest axons of each nerve in (d) (one way ANOVA followed by Tukey’s multiple comparisons test, *P*=0.0052; wild type: n=10 mice, Kdm6a KO/male: n=3 mice, Kdm6a KO/female: n=7). (g) Top: schematic time course of optic nerve regeneration model. Optic nerve regeneration was analyzed 2 weeks after the nerve crush, and the fluorescence (Alexa 594) tagged CTB was injected 2 days before collecting the optic nerve. Bottom: representative 2D projected confocal images of whole mount cleared optic nerves of *Kdm6a* mutant knockin (Kdm6a-mut KI) and wild type mice 2 weeks after ONC. The regenerating axons were labeled with CTB-Alexa 594. The red dotted lines indicate the nerve crush sites. The columns on the right display magnified images of the areas in white, dashed boxes on the left, showing axons at 0.25, 0.75, and 1.25 mm distal to the crush sites. Scale bar, 200 µm for the left column and 150 µm for the magnified images. (h) Quantification of regenerating axons at different distances distal to the nerve crush site (0.25 to 1.75 mm) in Kdm6a-mut KI and wild type mice after ONC. Bar graph showing robust optic nerve regeneration in Kdm6a- mut KI mice (two tailed student’s *t* tests, *P*<0.05; Wild type: n=5 mice, Kdm6a-mut KI: n=5 mice). (i) Quantification of the average length of the top 5 longest axons of each nerve in (g) (two tailed student’s *t* tests, *P*=0.0087; Wild type: n=5 mice, Kdm6a-mut KI: n=5 mice). (j) Representative images of flat whole mount retina immunostained for neuronal marker Tuj1 (green), indicating increased RGC survival rates in either male *Kdm6a* knockout (Kdm6a KO/male) or female *Kdm6a* knockout (Kdm6a KO/female) mice compared with that in wild type mice. Scale bar, 50 µm. (k) Quantification of RGC survival rates in wild type, male *Kdm6a* knockout or female *Kdm6a* knockout mice after ONC (one way ANOVA followed by Tukey’s multiple comparisons test, *P*=0.0291; n=3 mice for each condition). (l) Representative images of flat whole mount retina immunostained for neuronal marker Tuj1 (green), indicating increased RGC survival rates in female *Kdm6a* knockout (Kdm6a KO/female) mice compared with that in wild type mice after NMDA induced injury. Scale bar, 50 µm. (m) Quantification of RGC survival rate in (l) (two tailed student’s *t* tests, *P*=0.0485; n=3 mice for each condition). *, **, \*\*\**P* < 0.05, 0.01, 0.001, respectively. Data are represented as mean ±SEM. See also Extended Data Figs. 1–3.

We therefore obtained the *Kdm6a* floxed mice to examine its role in optic nerve regeneration. Because the *Kdm6a* gene is located on the X chromosome, the female mice have both alleles of *Kdm6a* floxed (*Kdm6a^f/f^*), whereas the male mice having only allele floxed with the *kdm6c* (*Uty*) gene on the Y chromosome (*Kdm6a^f/y^*). *Kdm6a* was knocked out by intravitreal injection of adeno-associated virus (AAV2) vector encoding the Cre recombinase. The transduction rate of AAV2-Cre in RGCs was 88.8 ± 7.56% (n=2 retinas), evaluated by co-immunostaining the whole mount retina with antibodies against Tuj1 and Cre (**Extended Data** Figure 3a). The mRNA level of *Kdm6a* in the purified RGCs from the *Kdm6a^f/f^*/AAV2-Cre mice was markedly downregulated compared with that from the control *Kdm6a^f/f^*/AAV2-GFP mice (**Extended Data** Figure 3b). In consistent, immunostaining analysis 2 weeks after viral infection showed significantly reduced level of Kdm6a in RGCs in AAV2-Cre injected retinas (**Extended Data** Figure 3c**, d**). As expected, deleting the demethylase Kdm6a resulted in significantly elevated level of H3K27me3 in RGCs (**Extended Data** Figure 3e**, f**).

To determine how knocking out *Kdm6a* in RGCs affect RGC survival and optic nerve regeneration, we performed ONC two weeks after the virus injection in *Kdm6a^f/f^*/AAV2-Cre (female), *Kdm6a^f/y^*/AAV2-Cre (male) or the control mice (wild type littermate mice injected with AAV2-Cre). Two weeks after ONC, RGC axons were labeled anterogradely by intravitreal injection of the Alexa-594-conjugated cholera toxin β (CTB) and optic nerve regeneration was assessed (**Figure 2d**). Fluorescence 3D images of tissue-cleared optic nerves were acquired with confocal microscopy as described in our previous studies.[11, 28, 58] Wild type mice showed minimal optic nerve regeneration passing the crush sites, whereas knocking out *Kdm6a* in female mice significantly promoted optic nerve regeneration (**Figure 2d-f**). Knocking out *Kdm6a* in male mice also induced enhanced optic nerve regeneration, but to a much lesser degree (**Figure 2d-f**), likely due to the presence of the *Kdm6c* gene on the Y chromosome with reduced demethylase activity.[59] In addition, we obtained Kdm6a enzyme-dead knockin mice (Kdm6a-mut KI) which possess the H1146A and E1148A point mutations in exon 24.[60] To investigate the role of Kdm6a enzymatic activity in optic nerve regeneration, we first performed ONC in either Kdm6a-mut KI mice or the control mice (wild type). Two weeks later, optic nerve regeneration was analyzed by quantifying the number of CTB-594 labeled axons (**Figure 2g**). The number and length of regenerating axons in Kdm6a-mut KI mice were enhanced significantly compared to those in wild type mice, providing further evidence that the histone demethylase activity of Kdm6a functions to repress optic nerve regeneration (**Figure 2g-i**). Next, we investigated whether Kdm6a affected the RGC survival with two widely used *in vivo* cell death models, including the ONC and the NMDA receptor-mediated excitotoxity, a shared pathological pathway of neurodegenerative diseases. The results showed that knocking out *Kdm6a* in either female or male mice significantly protected RGCs from cell death to the same extent (**Figure 2j, k**). These results indicated that Kdm6c on the Y chromosome was not involved in ONC-induced RGC death, different from that of axon regeneration. In addition, deleting *Kdm6a* also markedly increased RGC survival after NMDA treatment (**Figure 2l, m**), suggesting its wide range of neuroprotection after different types of neural injuries.

### 2.3. Knocking out *Kdm6b* enhances RGC survival, but poorly promotes axon regeneration

We next assessed the effects of deleting *Kdm6b* in adult RGCs on optic nerve regeneration. The expression pattern of Kdm6b within RGCs was firstly analyzed at different time points after ONC. The dot plot and violin plot obtained from scRNA-seq dataset[57] showed that the mRNA level of *Kdm6b* was increased upon ONC (**Extended Data** Figure 2a**, c**). By performing immunofluorescence staining, we further confirmed the upregulation of Kdm6b in RGCs after ONC (**Figure 3a-c**). In contrast to Kdm6a, the staining results showed that Kdm6b was mainly localized in the cytoplasm of RGCs before and after ONC, suggesting that it might not be the major histone demethylase in RGCs. Indeed, when Kdm6b in RGCs was knocked out with AAV2-Cre in *Kdm6b^f/f^* mice (**Extended Data** Figure 3g-i), the level of H3K27me3 in RGCs was actually slightly decreased but without statistical significance (**Extended Data** Figure 3j**, k**). When optic nerve regeneration was quantified, knocking out *Kdm6b* did not significantly increase the number of regenerating axons at different distances from the crush site (**Figure 3d, e**). Moreover, when both *Kdm6a* and *Kdm6b* were knocked out, there was no additive promoting effect on optic nerve regeneration (**Figure 2d and Figure 3d, f**) compared to Kdm6a knockout alone, further confirming that Kdm6b was not involved in regulating optic nerve regeneration. When RGC survival was examined, deleting Kdm6b significantly enhanced RGC survival, and double knocking out *Kdm6a/b* showed additive effect on RGC survival (**Figure 3g, h**). Together, these findings provided clear evidence that Kdm6b did not regulate optic nerve regeneration, in contrast to that of Kdm6a or that observed in sensory neurons. However, Kdm6b is able to regulate RGC survival but with different underlying mechanisms from that of Kdm6a.

**Fig. 3.**
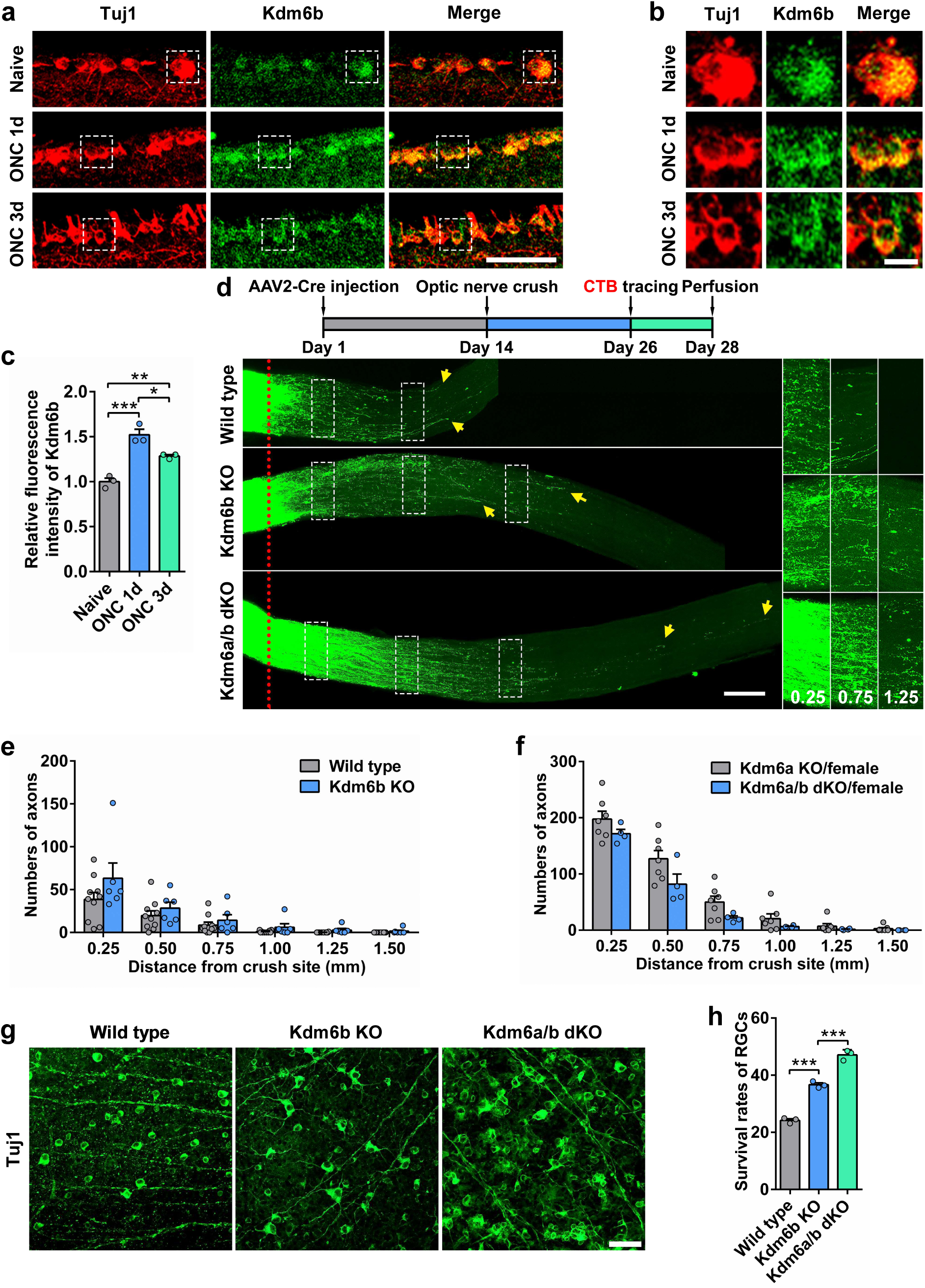
Knocking out H3K27 demethylase *Kdm6b* in mouse retinal ganglion cells (RGCs) markedly enhanced neuronal survival, but poorly promoted axon regeneration. (a) Representative images of Kdm6b immunostaining in retina sections showing increased level of Kdm6b in RGCs after ONC. Images were stained for neuronal marker Tuj1 (red) and Kdm6b (green). Scare bar, 50 μm. (b) Enlarged images of areas indicated by the dashed white boxes in (a). Scale bar, 10 µm. (c) Quantification of relative fluorescence intensity of Kdm6b immunostaining in (a) (one way ANOVA followed by Tukey’s multiple comparison test, P=0.0004; n=3 independent experiments for each condition). (d) Top: schematic time course of optic nerve regeneration model. Bottom: representative 2D projected confocal images of whole mount cleared optic nerves of wild type, *Kdm6b* knockout (Kdm6b KO), *Kdm6a* and *Kdm6b* double knockout (Kdm6a/b dKO) mice 2 weeks after ONC. The regenerating axons were labeled with CTB-Alexa 594. Yellow arrows indicate the distal ends of regenerating axons. The red dotted lines indicate the nerve crush sites. The columns on the right display magnified images of the areas in white, dashed boxes on the left, showing axons at 0.25, 0.75, and 1.25 mm distal to the crush sites. Scale bar, 200 µm for the left column and 100 µm for the magnified images. (e) Quantification of regenerating axons at different distances distal to the nerve crush site (0.25 to 1.50 mm) in wild type or *Kdm6b* knockout mice (Kdm6b KO) after ONC. Bar graph showing no difference in axon regeneration between wild type and Kdm6b KO (two tailed student’s *t* tests, *P*>0.05; wild type: n=10 mice, Kdm6b KO: n=6 mice). (f) Quantification of regenerating axons at different distances distal to the nerve crush site (0.25 to 1.50 mm) in female *Kdm6a* knockout (Kdm6a KO/female) or female *Kdm6a*/*Kdm6b* double knockout mice (Kdm6a/b dKO/female) after ONC. Bar graph showing no additive/synergistic effects of *Kdm6a*/*Kdm6b* co-deletion on optic nerve regeneration (two tailed student’s *t* tests, *P*>0.05; Kdm6a KO: n=7 mice, Kdm6a/b dKO: n=4 mice). (g) Representative images of flat whole mount retina immunostained for neuronal marker Tuj1 (green), indicating increased RGC survival rates in either *Kdm6b* knockout (Kdm6b KO) or *Kdm6a*/*Kdm6b* double knockout (Kdm6a/b dKO) mice compared with that in wild type mice. Co-deleting *Kdm6a*/*Kdm6b* resulted in additive/synergistic effects on RGC survival. Scale bar, 50 µm. (h) Quantification of RGC survival rate in (g) (one way ANOVA followed by Tukey’s multiple comparisons test, *P*<0.0001; n=3 mice for each condition). *, **, \*\*\**P* < 0.05, 0.01, 0.001, respectively. Data are represented as mean ±SEM. See also Extended Data Figs. 2, 3.

### 2.4. Pharmacological inhibition of Kdm6a/b in RGCs after ONC still promotes optic nerve regeneration and RGC survival

To explore the potential of translational application, we tested whether a delayed inhibition of Kdm6a in RGCs after the injury could also promote optic nerve regeneration. We used GSK-J4, a small molecule catalytic site inhibitor selective for the H3K27me3-specific demethylase subfamily (*Kdm6a/b*).[61] GSK-J4 was injected intravitreally every 3 days for 2 weeks after the ONC. The immunostaining results showed that GSK-J4 treatment significantly increased H3K27me3 levels in RGCs, indicating the inhibitory effect on Kdm6a/b (**Figure 4a, b**). When axon regeneration was examined, the results showed that in control mice DMSO had little effect on optic nerve regeneration, whereas significant regeneration was observed in mice with GSK-J4 treatment (**Figure 4c, d**). The average lengths of the top 5 longest regenerating axons in mice injected with GSK-J4 were markedly longer than those in control mice with DMSO injection (**Figure 4e**). Delayed pharmacological inhibition of Kdm6a also promoted RGC survival (**Figure 4f, g**). Collectively, these results indicated that inhibiting the demethylase activity of Kdm6a after neural injury was able to effectively enhance CNS axon regeneration and neuronal survival.

**Fig. 4.**
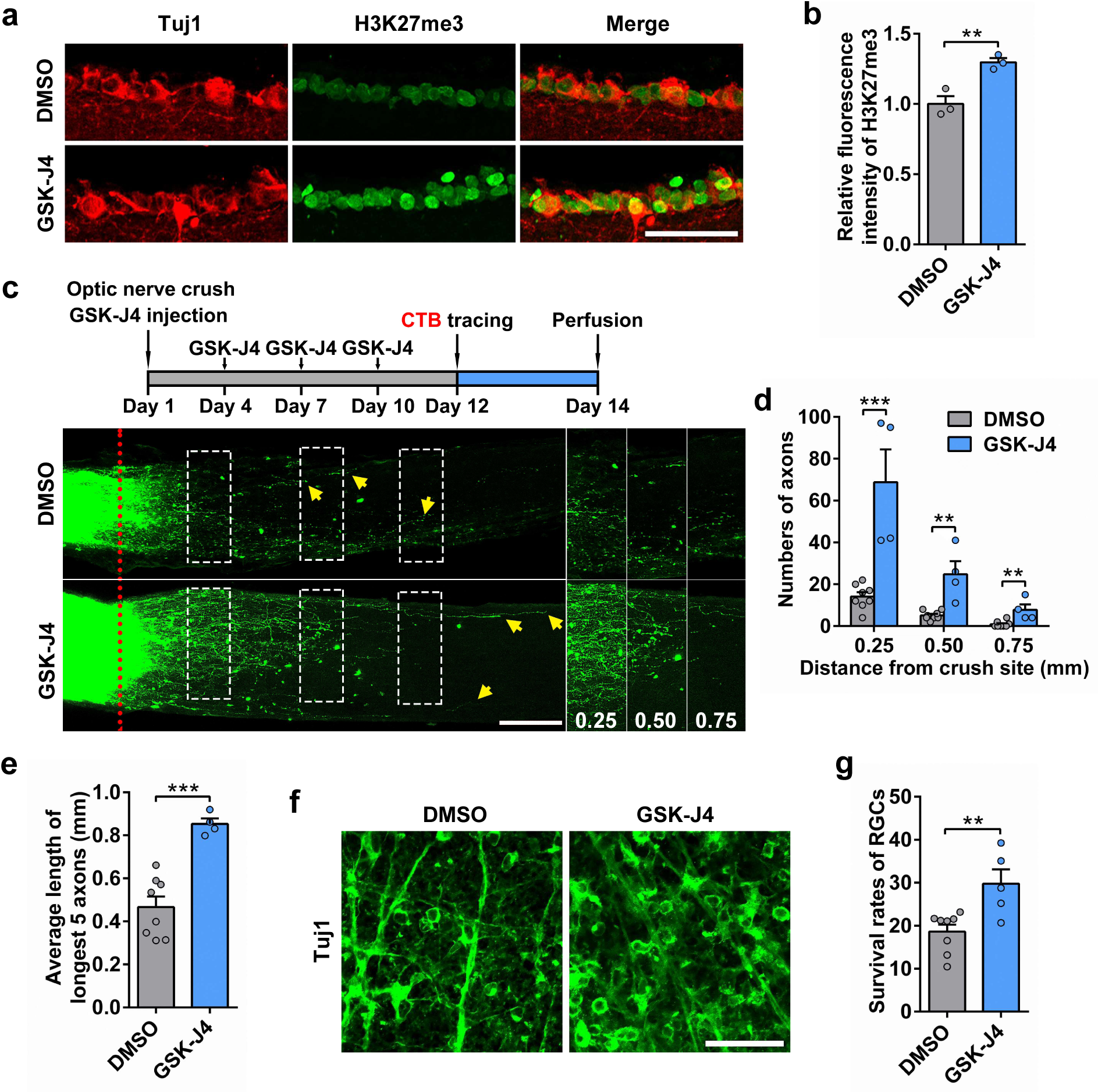
**Pharmacological delayed inhibition of Kdm6a/b in mouse retinal ganglion cells (RGCs) also promoted optic nerve regeneration and RGC survival.** (a) Confocal images of retinal sections from wild type mice injected with Kdm6a/b inhibitor GSK-J4 or DMSO (as control) showing significantly increased level of H3K27me3 in RGCs by blockade of histone demethylases activity. The sections were stained for neuronal marker Tuj1 (red), which labeled RGCs, and H3K27me3 (green). Scale bar, 50 µm. (b) Quantification of relative fluorescence intensity of H3K27me3 immunostaining shown in (a) (two tailed student’s *t* tests, *P*=0.0091; n=3 mice for each condition). (c) Top: schematic time course of optic nerve regeneration model. Specifically, GSK-J4 or DMSO was injected once every 3 days for 4 times after ONC. Bottom: representative 2D projected confocal images of whole mount cleared optic nerves of wild type mice with intravitreal injection of GSK-J4 or DMSO 2 weeks after ONC. The regenerating axons were labeled with CTB-Alexa 594. The red dotted lines indicate the nerve crush sites. The columns on the right display magnified images of the areas in white, dashed boxes on the left, showing axons at 0.25, 0.50, and 0.75 mm distal to the crush sites. Scale bar, 200 µm for the left column and 130 µm for the magnified images. (d) Quantification of regenerating axons at different distances distal to the nerve crush site (0.25 to 0.75 mm) in injected GSK-J4 or DMSO mice after ONC. Bar graph showing robust optic nerve regeneration induced by GSK- J4 (two tailed student’s *t* tests, *P*<0.01; DMSO: n=8 mice, GSK-J4: n=4 mice). (e) Quantification of the average length of the top 5 longest axons of each nerve in (c) (two tailed student’s *t* tests, *P*=0.0004; DMSO: n=8 mice, GSK-J4: n=4 mice). (f) Representative images of flat whole mount retina immunostained for neuronal marker Tuj1 (green), indicating increased RGC survival rates in injected GSK-J4 mice compared with that in mice with DMSO injection. Scale bar, 50 µm. (g) Quantification of RGC survival rate in (f) (two tailed student’s *t* tests, *P*=0.0063; DMSO: n=8 mice, GSK-J4: n=5 mice). **, \*\*\**P* < 0.01, 0.001, respectively. Data are represented as mean ± SEM.

### 2.5. *Kdm6a* knockout enhances optic nerve regeneration via distinct pathways from that of Pten deletion

Activation of the mammalian target of rapamycin (mTOR) signaling in adult RGCs is one of the major pathways for promoting optic nerve axon regeneration downstream of several genes, such as Pten, Lin28, Osteopontin, and Akt/GSK3, etc.[62] Thus, we investigated if Kdm6a deletion also enhanced optic nerve regeneration through the same pathway. Immunostaining of retina sections showed that at 3 days after injury, deleting *Kdm6a* alone in RGCs had no effect on the percentage of phospho-S6 (p-S6) positive RGCs (**Figure 5a, b**), indicating that the mTOR pathway was not activated. In contrast, Kdm6a/Pten co-deletion dramatically increased the percentage of p-S6+ RGCs (**Figure 5c, d**), as well as the level of H3K27me3 in RGCs (**Figure 5e, f**). These results indicated that *Kdm6a* knockout led to novel downstream signaling pathways distinct from that of Pten deletion and mTOR activation. We therefore examined the effects of co-deleting *Kdm6a* and *Pten* in adult RGCs on optic nerve regeneration. AAV2-Cre were injected into the vitreous body of *Pten^f/f^* or *Pten^f/f^/Kdm6a^f/f^* mice to delete *Pten* or *Pten*/*Kdm6a*. Real-time PCR analysis showed that the mRNA levels of both *Kdm6a* and *Pten* were significantly reduced in *Pten*/*Kdm6a* double knockout mice (**Extended Data** Figure 4a). Optic nerve regeneration was examined 2 weeks after the ONC. The results showed that concomitant deletion of Kdm6a and Pten in adult RGCs triggered faster optic nerve regeneration and more regenerative axons, compared with Pten deletion alone (**Figure 5g-i**). At 2 weeks after ONC, some long regenerative axons almost reached the region proximal to the optic chiasm (**Figure 5g**). Furthermore, the additive effects of *Kdm6a/Pten* double knockout were persistent to 6 weeks after ONC (**Extended Data** Figure 4b-d). In *Pten* single knockout mice almost no regenerating axons could be identified in the proximal end of the optic chiasm, whereas some regenerating axons in *Kdm6a*/*Pten* double knockout mice reached and crossed the optic nerve-chiasm transition zone (OCTZ) (**Extended Data** Figure 4b). To evaluate whether regenerating axons entered the optic chiasm and continued to grow into the optic tracts, we performed a distal intra-orbital optic nerve crush that shortened the distance axons needed to grow to reach the optic chiasm (**Figure 5j**). At 4 weeks after the injury, in *Kdm6a*/*Pten* double knockout mice many regenerating axons entered the chiasm, whereas in significant contrast almost no axons in *Pten* knockout mice reached the chiasm (**Figure 5j, k**). Together, these results demonstrated that co-deletion of Kdm6a and Pten could result in an additive effect on promoting long-distance axon regeneration.

**Fig. 5.**
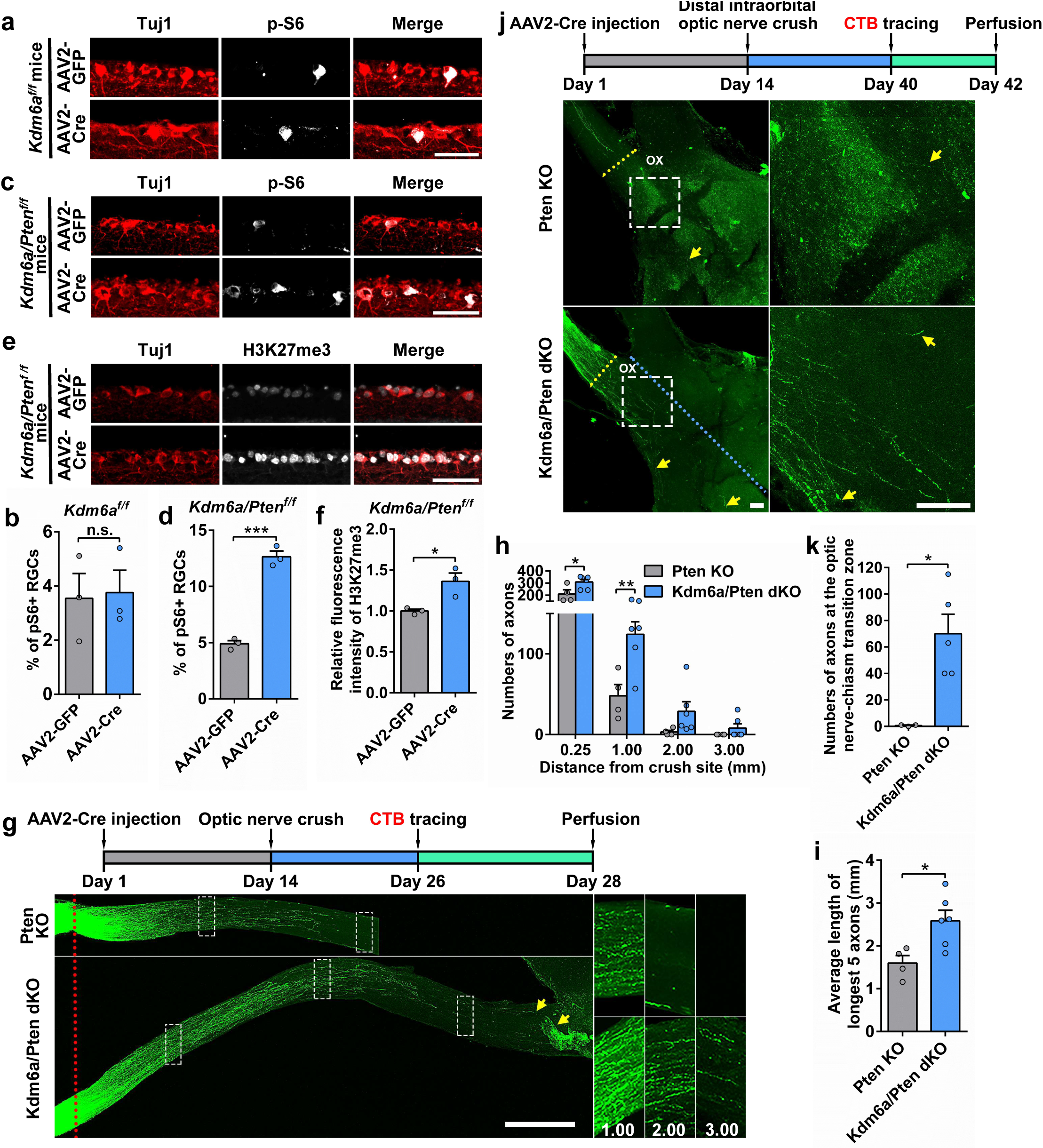
Synergistic promoting effects of *Kdm6a* and *Pten* double knockout on axon regeneration after ONC. (a) Confocal images of retinal sections from *Kdm6a^f/f^* mice injected with of AAV2-GFP or AAV2-Cre showing no effect of *Kdm6a* single deletion on the percentage of p-S6+ RGCs in the ganglion cell layer (GCL). The sections were stained for neuronal marker Tuj1 (red), which labeled RGCs, and p-S6 (gray). Scale bar, 50 µm. (b) Quantification of p-S6+ RGCs in (a) (two tailed student’s *t* tests, *P*=0.8747; n=3 mice for each condition). (c) Confocal images of retinal sections from *Kdm6a/Pten^f/f^* mice injected with of AAV2-GFP or AAV2-Cre showing markedly increased percentage of p-S6+ RGCs in the ganglion cell layer (GCL) via Cre-mediated double deletion. The sections were stained for neuronal marker Tuj1 (red), which labeled RGCs, and p-S6 (gray). Scale bar, 50 µm. (d) Quantification of p-S6+ RGCs in (c) (two tailed student’s *t* tests, *P*=0.0002; n=3 mice for each condition). (e) Confocal images of retinal sections from *Kdm6a/Pten^f/f^* mice injected with of AAV2-GFP or AAV2-Cre showing significantly increased level of H3K27me3 in RGCs via Cre-mediated double deletion. The sections were stained for neuronal marker Tuj1 (red), which labeled RGCs, and H3K27me3 (gray). Scale bar, 50 µm. (f) Quantification of relative fluorescence intensity of H3K27me3 immunostaining shown in (e) (two tailed student’s *t* tests, *P*=0.0252; n=3 mice for each condition). (g) Top: schematic time course of optic nerve regeneration model. Bottom: representative 2D projected confocal images of whole mount cleared optic nerves of *Pten* knockout (Pten KO) or *Kdm6a*/*Pten* double knockout (Kdm6a/Pten dKO) mice 2 weeks after ONC. The regenerating axons were labeled with CTB-Alexa 594. Yellow arrows indicate the distal ends of regenerating axons. The red dotted lines indicate the nerve crush sites. The columns on the right display magnified images of the areas in white, dashed boxes on the left, showing axons at 1.00, 2.00, and 3.00 mm distal to the crush sites. Scale bar, 500 µm for the left column and 250 µm for the magnified images. (h) Quantification of regenerating axons at different distances distal to the nerve crush site (0.25 to 3.00 mm) in *Pten* knockout (Pten KO) or *Kdm6a*/*Pten* double knockout mice (Kdm6a/Pten dKO) 2 weeks after ONC (two tailed student’s *t* tests, *P<*0.05; Pten KO: n=4 mice, Kdm6a/Pten dKO: n=6 mice). (i) Quantification of the average length of the top 5 longest axons of each nerve in (g) (two tailed student’s *t* tests, *P*=0.0184; Pten KO: n=4 mice, Kdm6a/Pten dKO: n=6 mice). (j) Top: schematic time course of optic nerve regeneration model. The optic nerve was crushed at the distal intraorbital site. Optic nerve regeneration was analyzed 4 weeks after the nerve crush. Bottom: Representative 2D projected confocal images of whole mount cleared optic chiasm areas of *Pten* knockout (Pten KO) or *Kdm6a/Pten* double knockout (Kdm6a/Pten dKO) mice 4 weeks after distal intraorbital optic nerve crush. The right column shows enlarged images of areas indicated by the dashed white boxes in the left column. The regenerating axons at the optic chiasm were labeled with CTB-Alexa 594. Yellow arrows indicate the distal ends of regenerating axons. Blue and yellow dotted lines indicate the chiasm midline and the optic nerve-chiasm transition zone, respectively. OX indicates the optic chiasm. Scale bar, 100 µm. (k) Quantification of regenerating axons reaching the optic chiasm in (j) (two tailed student’s *t* tests, *P*=0.0125; Pten KO: n=3 mice, Kdm6a/Pten dKO: n=5 mice). n.s. *P* > 0.05; *, **, \*\*\**P* < 0.05, 0.01, 0.001, respectively. Data are represented as mean ± SEM. See also Extended Data Fig. 4.

### 2.6. Regenerating axons show complex growth patterns in *Kdm6a* or *Kdm6a*/*Pten* knockout mice

Tissue clearing and 3D imaging allowed us to perform detailed analyses of axonal tip morphology and axon trajectory at the single axon level, revealing potential cellular mechanism associated with axon regeneration. As described in our previous study,[58] we first quantified the sizes of the distal axon ends (growth cones) of each regenerating axon. We found that Kdm6a deletion significantly reduced the sizes of the distal axon ends, compared to wild type (**Figure 6a-c**). Specifically, in wild type mice, the majority of weakly regenerated short axons had bulbous structures at the distal tips. About 20% of the axons formed the bigger terminal swellings (tip/shaft ratio > 4) defined as retraction bulb-like structures, which are the characteristic structures of non-regenerating axons (**Figure 6c**). In contrast, most regenerating axons in male or female mice with Kdm6a deletion showed slim growth cones (**Figure 6a-c**), leading to enhanced axon extension. Furthermore, detailed analysis of axon trajectory showed that in either wild type or *Kdm6a* knockout mice, within the optic nerve regenerating axons often had a meandering path and many of them made U-turns (**Figure 6a, d**), indicating the inhibitory nature of the mature visual system. Interestingly, deleting *Kdm6a* seemed to enhance the U-turn rate, likely due to increased number of regenerating axons.

**Fig. 6.**
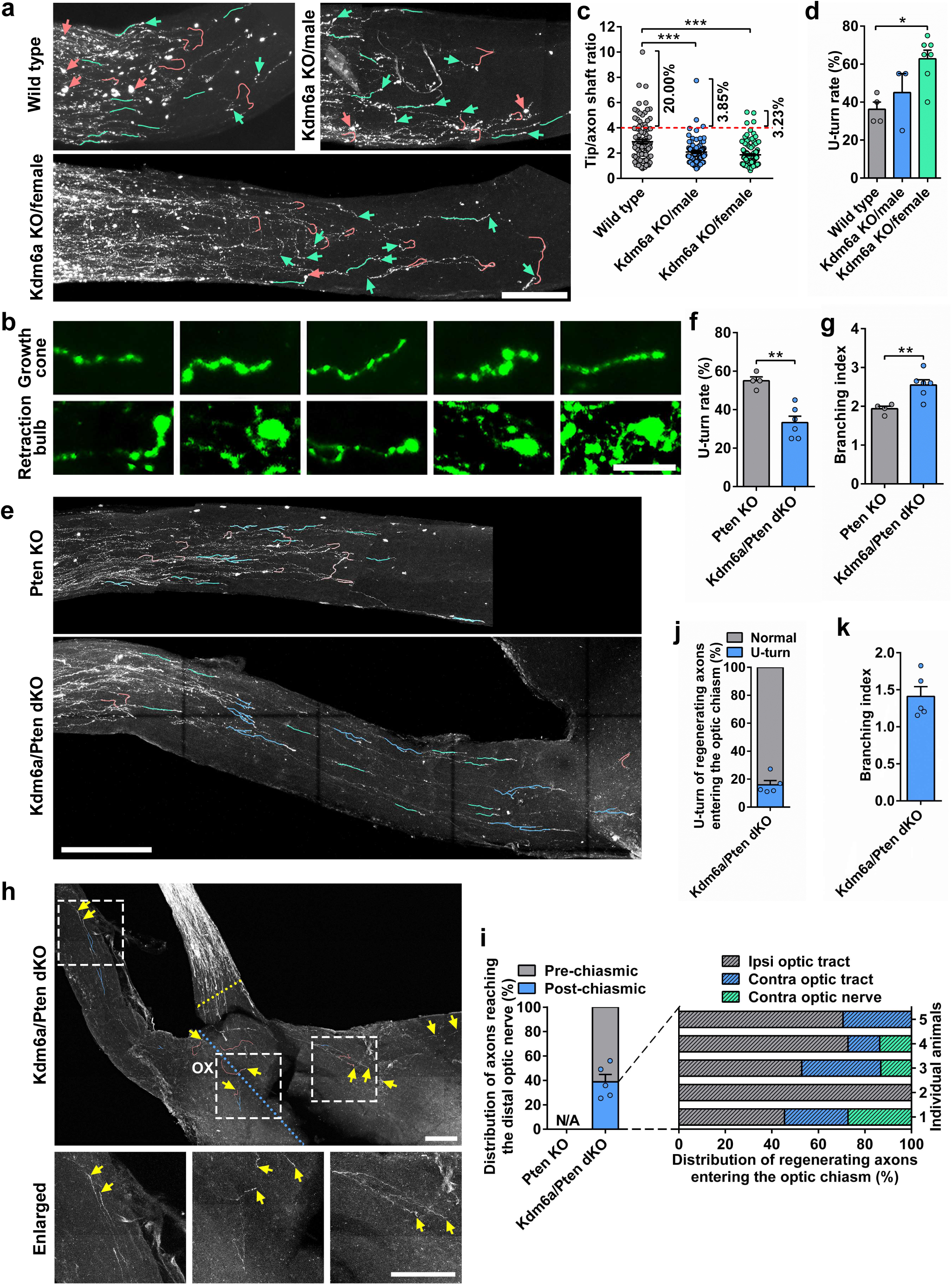
*Kdm6a* or *Kdm6a*/*Pten* knockout induced changes of axonal morphology. (a) Representative images of optic nerves showing that *Kdm6a* deletion modified RGC axonal morphology 2 weeks after ONC. Neurite tracing of single axon showing straight trajectories (green lines) and U-turns (red lines). Red and green arrows indicate retraction bulbs and growth cones, respectively. Scale bar, 200 μm. (b) Representative images of growth cones and retraction bulbs found in different optic nerves. Scale bar, 25 μm. (c) Quantification of the size of retraction bulbs and growth cones by tip/axon shaft ratio in (a) (one way ANOVA followed by Tukey’s multiple comparisons test, *P*<0.0001; n=105, 78, and 124 axons pooled from 7 mice for wild type, 3 mice for Kdm6a KO/male, and 7 mice for Kdm6a KO/female, respectively). (d) Quantification of U-turn rate in (a) (one way ANOVA followed by Tukey’s multiple comparisons test, *P=*0.0126; wild type: n=4 mice, Kdm6a KO/male: n=3 mice, Kdm6a KO/female: n=7; top 20 longest axons were analyzed for each mouse). (e) Representative images of U-turns and branches found in the distal nerve region. Neurite tracing showing that co-deletion of *Kdm6a*/*Pten* in RGCs led to markedly decreased U-turns and increased branches compared to *Pten* deletion alone. Green lines, red lines and blue lines indicate straight trajectories, U-turns and branches, respectively. Scale bar, 200 μm. (f) Quantification of U-turn rate in (e) (two tailed student’s *t* tests, *P*=0.0013; Pten KO: n=4 mice, Kdm6a/ Pten dKO: n=6 mice; top 20 longest axons were analyzed for each mouse). (g) Quantification of branching index by tip number/axon number (the top 20 longest axons) ratio in (e) (two tailed student’s *t* tests, *P*=0.0088; Pten KO: n=4 mice, Kdm6a/Pten dKO: n=6 mice). (h) Neurite tracing of regenerating axon into the optic chiasm showing complex growth patterns in *Kdm6a/ Pten* double knockout mice. The bottom row shows enlarged images of areas indicated by the dashed white boxes in the top row. The regenerating axons at the optic chiasm were labeled with CTB-Alexa 594. Yellow arrows indicate the distal ends of regenerating axons. Blue and yellow dotted lines indicate the chiasm midline and the optic nerve-chiasm transition zone, respectively. Green lines, red lines and blue lines indicate straight trajectories, U-turns and branches, respectively. OX indicates the optic chiasm. Scale bar, 200 µm. (i) Distribution of regenerating axons reaching the distal optic nerve (Pre-chiasmic) and entering the optic chiasm (Post-chiasmic) in (h). The dashed lines showing quantification of post-chiasmic axons projecting into the ipsilateral optic tract (Ipsi optic tract), contralateral optic tract (Contra optic tract), and contralateral optic nerve (Contra optic nerve) in 5 individual mice. N/A, not applicable. (j) Quantification of U-turn rate in (h) (n=5 mice; all axons into the optic chiasm were analyzed for each mouse). (k) Quantification of branching index by tip number/axon number (regenerating axons into the optic chiasm) ratio in (h) (n=5 mice). *, **, \*\*\**P* < 0.05, 0.01, 0.001, respectively. Data are represented as mean ±SEM. See also Extended Data Fig. 4.

Based on whole tissue 3D imaging, we also tracked the trajectories of individual RGC axons in the distal region of the optic nerve (**Figure 6e**). Within the distal optic nerve, many regenerating RGC axons induced by Pten deletion alone showed wandering trajectories with frequent U-turns (**Figure 6f**), similar to that of *Kdm6a* knockout alone (**Figure 6d**). However, the U-turn rate was markedly reduced when *Kdm6a* and *Pten* were both knocked out (**Figure 6f and Extended Data** Figure 4e). Moreover, we noticed that some regenerating RGC axons extended multiple branches from the axonal shafts or at the axon ends (**Figure 6e**). Quantification of axonal branching showed that compared to Pten deletion alone, the combination of Kdm6a and Pten deletion led to increased formation of axon branches (**Figure 6g**), indicating enhanced axon growth capacity.

To better examine how regenerating axons behaved at the optic chiasm, we analyzed morphological characteristics of regenerating axons passing the optic chiasm in the distal intra- orbital optic nerve crush mice with Kdm6a/Pten co-deletion (**Figure 6h**). We observed that in the mice with Kdm6a/Pten co-deletion, more than 60% of regenerating axons stopped growing or turned back at the entry area of the chiasm, indicating the optic chiasm as a significant inhibitory barrier (**Figure 6h, i**). For regenerating axons entering the chiasm most of them were found in the ipsilateral optic tract (**Figure 6h, i**), indicating axon misguidance. A few regenerating axons crossed the midline into the contralateral optic tract or extended into the opposite uninjured optic nerve (**Figure 6h, i**). We also assessed growth patterns of RGC axons entering the chiasm in *Kdm6a*/*Pten* double knockout mice (**Figure 6h**). The results showed that within the optic chiasm, approximately 20% of axons made sharp turns, and axonal branches were observed in different regions (**Figure 6h, j, k**). Notably, RGC axons entering the chiasm showed reduced U-turns and axon branching (**Figure 6j, k**) compared with those within optic nerve (**Figure 6f, g**), suggesting spatial differences of the environment between the two regions.

### 2.7. Kdm6a deletion in RGCs triggers activation of developmental-like transcriptomic programs

To further gain mechanistic insights into promoting effect of Kdm6a on axon regeneration, we performed RNA sequencing (RNA-seq) of the enriched RGCs across different developmental stages (postnatal day 1, 14 and 21) and at 3 days after injury of adult (P42) wild type (AAV2-GFP) or Kdm6a knockout mice (AAV2-Cre). Dissociated RGCs were labeled with Thy1.2 antibody, subjected to fluorescence-activated cell sorting (FACS) for RGC enrichment, and identified by labelling for Tuj1 (**Extended Data** Figure 5). Two RNA-seq datasets (13 bulk RNA-seq libraries) were generated from FACS-purified RGCs (**Extended Data** Figure 6a). It is widely believed that during neuronal maturation the changes in cellular states at the transcriptional level underlie their reduced ability to support regeneration. Young neurons have strong axon growth ability to form neural circuits, whereas mature neurons lose the ability to grow and change to support stable synaptic function.[63] Significant differentially expressed genes (DEGs) were first recognized by comparing the gene expression profiles between P1 and P14 RGCs, but modest DEGs between P14 and P21 RGCs (**Extended Data** Figure 6b**, c and e**). These results indicated that RGCs changed their cellular states from young to mature between P1 and P14. Gene enrichment ontology (GO) analysis showed that downregulated DEGs in P14 RGCs compared to P1 RGCs were associated with axonogenesis, developmental cell growth and neuron projection guidance, supporting that axon growth ability reduced during neuronal maturation (**Extended Data** Figure 6d**, f**). We further compared the transcriptomic profiles of injured adult control or *Kdm6a* knockout RGCs (P42) to RGC maturational datasets. Hierarchical clustering analysis (HCA) revealed that the profiles were grouped into two main clusters with similar gene expression patterns (**Extended Data** Figure 7a). The first cluster included P1, P14 and AAV2-Cre (*Kdm6a* knockout P42 RGCs), while the second cluster involved P21 and AAV2-GFP (control P42 RGCs). In addition, the transcriptomic profiles of injured *Kdm6a* knockout RGCs (AAV2-Cre) were more similar to the P1 and P14 than to wild type control RGCs (AAV2-GFP) and P21 RGCs. Furthermore, principal component analysis (PCA), the linear dimensionality reduction method, was adopted for precisely screening the significantly components and finding potential clusters across all samples. The results showed that the first principal component (Dim 1) contribute the most (60.6%) for explaining the variance (**Figure 7a**). *Kdm6a* knockout (AAV2-Cre) clustered close to P1 along Dim 1 compared to wild type (AAV2-GFP), suggesting the strong commonalities of *Kdm6a* knockout and developmental stage RGCs (**Figure 7a**). Next, we identified genes with significant contribution in the PCA clustering, especially relevant to Dim 1. GO enrichment analysis was conducted to reveal the biological roles of genes involved in Dim 1 and 2. As shown in **Figure 7b**, Dim 1 explained most variance in the transcriptomic pattern of genes involved in postsynaptic organization, dendrite development, neuron projection organization, and neuron apoptotic process etc.

**Fig. 7.**
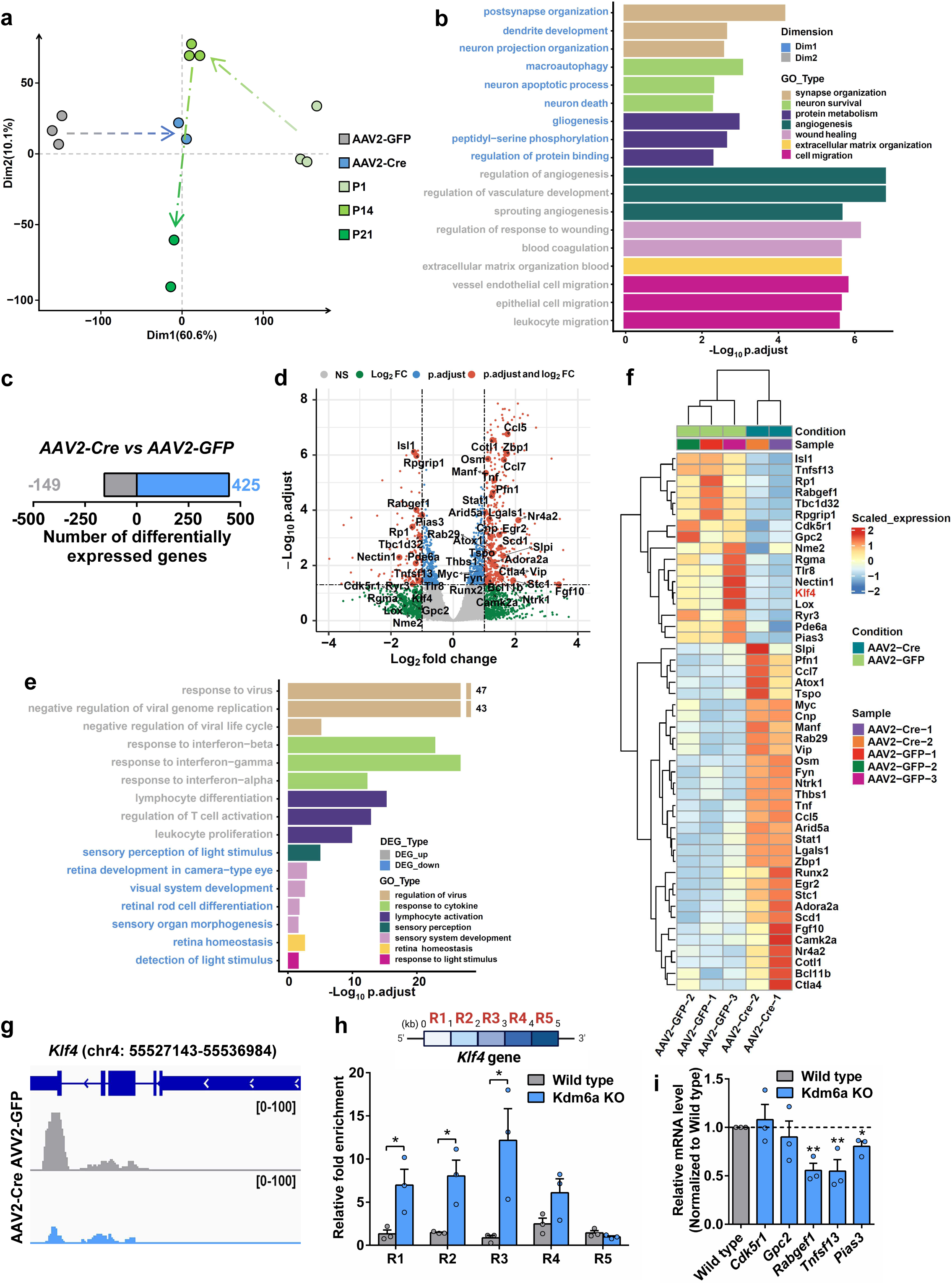
Deleting *Kdm6a* switched the RGC transcriptomics into a developmental-like state. (a) Principal component analysis (PCA) of transcriptomic profiles indicating the variation among all samples. Score plot shows a clear separation between P1, P14, P21, wild type (AAV2-GFP) and Kdm6a KO (AAV2-Cre). The first two dimensions (Dim) accounted for 70.7% of variance. Note that Dim 1 predominantly reflects differences between wild type and Kdm6a KO. (b) Selected Gene Ontology (GO) terms were generated separately for Dim 1 and 2 genes (adjusted *P*-value < 0.05). (c) Number of differentially expressed genes between wild type (AAV2-GFP) and Kdm6a KO (AAV2-Cre) at the threshold of absolute log_2_ Fold change > 1 and adjusted *P*-value < 0.05. (d) Volcano plot showing differential gene expression between wild type (AAV2-GFP) and Kdm6a KO (AAV2-Cre) RGCs. Positive or negative log_2_ Fold change indicate upregulation or downregulation in Kdm6a KO RGCs relative to wild type RGCs, respectively. Gray points (NS) indicate the genes with no significant change. Green points (Log_2_ FC) indicate the genes with absolute log_2_ Fold change > 1. Blue points (p.adjust) indicate the genes with adjusted *P*-value < 0.05. Red points (p.adjust and log_2_ FC) indicate that the genes are considered significantly different if absolute log_2_ Fold change > 1 and adjusted *P*-value < 0.05; vertical and horizontal reference lines at respective values. (e) Selected GO terms were generated separately for genes upregulated or downregulated (adjusted *P*- value < 0.05) by Kdm6a KO RGCs relative to wild type RGCs. (f) Heatmap of selected genes (absolute log_2_ Fold change > 1) relevant to axon regeneration and neuroprotection. (g) Sample genome tracks showing reduced expression levels of *Klf4* (chromosome 4 (chr4): 55527143-55536984) in the purified RGCs of *Kdm6a^f/f^* mice injected with AAV2-Cre, compared to *Kdm6a^f/f^* mice with AAV2-GFP injection. y axis indicates normalized reads. (h) Top: Schematic drawing of the 5 kilobase (kb) genomic regions (R1-R5) after the transcription start site (TSS) of the *Klf4* gene on chr4 were assayed in CUT&Tag-qPCR experiment using the antibody against H3K27me3 in the purified RGCs. In the H3K27me3 CUT&Tag-qPCR experiment, the genomic region R1-R5 were amplified from the purified RGCs to bind to H3K27me3. Bottom: CUT&Tag-qPCR results showing that deletion of Kdm6a (Kdm6a KO) led to increased interaction with R1, R2 and R3 regions of the *Klf4* gene (two tailed student’s *t* tests, R1: *P*=0.0402; R2: *P*=0.0240; R3: *P*=0.0378; n=3 independent experiments for each condition). (i) Real-time PCR results showing the expression of downregulated genes in RGCs from *Kdm6a* knockout mice (Kdm6a KO) after ONC (one sample *t* test, *Rabgef1*: *P*=0.0019; *Tnfsf13*: *P*=0.0095; *Pias3*: *P*=0.0111; n=3 independent experiments for each condition). *, \*\**P* < 0.05, 0.01, respectively. Data are represented as mean ±SEM. See also Extended Data Figs. 5-8.

By calculating the z-score of genes contributed to the top 20 enriched GO terms, it showed distinct heterogeneity between wild type and other groups, but significant homogeneity between Kdm6a deletion and P14 (**Extended Data** Figure 7b). Meanwhile, Double hierarchical clustered heatmap showed that many of these genes with top contribution in Dim 1 were associated with activation of developmental and growth programs. For instance, several genes, including Ehmt2,[64] Brsk1,[65] Eef1a2,[66] and Lrfn4,[67] have previously been shown to induce axon initiation, formation, sprouting and outgrowth (**Extended Data** Figure 7c). Thus, this finding suggests that knocking out *Kdm6a* in adult injured RGC triggers reactivation of developmental-like growth programs, leading to an immature cellular state.

To further identify gene regulatory networks that control the changes mediated by Kdm6a deletion, we performed a comparative analysis that profiled transcription in enriched RGCs. In total, 574 common DEGs were identified (425 significantly upregulated and 149 downregulated) at the thresholds of |log2 FC| > 1 (FC, fold change) and adjusted *P*-value of < 0.05 (**Figure 7c, d**). Furthermore, GO enrichment analysis indicated that the majority of genes downregulated by Kdm6a deletion were associated with sensory perception, sensory system development, retina homeostasis and response to light stimulus (**Figure 7e**), most of which related to mature neuronal functions. On the other hand, the highly enriched GO terms of up- regulated DEGs were mainly related to regulation of virus, response to cytokine and lymphocyte activation (**Figure 7e**) that were mostly non-neuronal genes. Several significantly altered genes were involved in axon regeneration (**Figure 7f**). For example, Klf4,[68] Myc,[9] and Thbs1[69] have been shown to play crucial roles in controlling axon regeneration. Additionally, Camk2 reactivation can protect RGCs in multiple injury/disease models.[70] Taken together, these results suggested that Kdm6a deletion promoted optic nerve axon regeneration via activating developmental-like growth programs and repressing mature neuron enriched genes.

### 2.8. Klf4 acts downstream of Kdm6a to regulate axon regeneration

H3K27me3 is well known to suppress gene expression. Here we showed that in sensory neurons upon peripheral axotomy the decreased level of Kdm6a/b and the subsequent elevated level of H3K27me3 acted to support the robust spontaneous sensory axon regeneration. Similarly, deletion of Kdm6a in RGCs resulted in an elevated level of H3K27me3, which in turn enhanced optic nerve regeneration. Therefore, it is very likely that the genes suppressed by H3K27me3 during axon regeneration function to repress axon regeneration.[71] In cultured RGCs, several Klf family proteins have been shown to suppress axon growth, including Klf1, 2, 4, 5, 9, 13, 14, 15, 16, among which Klf4 is well-known to repress axon regeneration *in vivo*.[4]

When Klf4 expression was examined in RGCs after *Kdm6a* knockout, the results showed that it was markedly downregulated (**Fig. 7f, g**). To investigate if *Klf4* was directly targeted by *Kdm6a* knockout-induced H3K27me3, we performed CUT&Tag with real-time PCR (CUT&Tag-qPCR) analysis in enriched RGCs using anti-H3K27me3 antibody for selected regions at 0 to 5000 bp after the transcription start site (TSS) of the *Klf4* gene on chromosome 4 (**Fig. 7h**). The results revealed that comparing with wild type RGCs, in RGCs lacking Kdm6a the interactions between H3K27me3 and 4 specific regions of the *Klf4* gene (R1-4) were significantly increased, especially at R1, R2 and R3 regions (**Fig. 7h**), indicating that in RGCs Klf4 expression was suppressed by Kdm6a-mediated H3K27me3 elevation.

Because Klf4 has also been shown to inhibit sensory axon regeneration *in vivo*,[72] we thus investigated if H3K27me3 could directly target *Klf4* and suppress its expression in sensory neurons during axon regeneration. By using an available single-cell RNA-seq (scRNA-seq) dataset of adult mouse DRG neurons,[73] we first examined the expression levels of all Klf family members (*Klf1-18*) in adult sensory neurons after SNI (**Extended Data** Figure 8a). The sequencing data suggested that at day 3 post-SNI, *Klf3, 4, 5, 7, 8, 9, 11, 12, 13, 14, 16* were reduced, whereas *Klf1, 2, 6, 10, 15, 17* were elevated. Additionally, *Klf18* was not expressed in adult sensory neurons. We then reprofiled the expression levels of all Klf family members in sensory neurons after *in vivo* axotomy by real-time PCR and detected transcripts for 17 (*Klf1- 17*) (**Extended Data** Figure 8b). We found that the mRNA levels of *Klf4* and *Klf5* were significantly downregulated in adult sensory neurons during peripheral nerve injury-induced axon regeneration (**Extended Data** Figure 8b), consistent with their roles in repressing axon growth. Furthermore, chromatin immunoprecipitation with PCR (ChIP-PCR) results showed that similar to that in RGCs, tri-methylated H3K27 interacted with 2 specific regions (R2 and R3) of the *Klf4* gene, indicating that H3K27me3 directly bind to the promoter region of *Klf4* in adult sensory neurons (**Extended Data** Figure 8c). Importantly, by using *in vivo* electroporation, we found that overexpressing Klf4 in adult sensory neurons significantly blocked axon regeneration *in vivo* (**Extended Data** Figure 8d**, e**), supporting its functional roles in repressing axon regeneration. Together, these results provided clear evidence that Kdm6a-H3K27me3 signaling was able to enhance mammalian axon regeneration by suppressing the regeneration repressor Klf4. In addition to Klf4, by examining downregulated genes in RGCs from *Kdm6a* knockout mice (**Fig. 7f**) with real-time PCR analysis, several other axon growth repressors were identified as potential downstream targets suppressed by the Kdm6a-H3K27me3 signaling, including *Rabgef1*, *Tnfsf13* and *Pias3* (**Fig. 7i**).

## 3. Discussion

Recent progresses in axon regeneration research have shown that significant changes in gene transcription underlie the regenerative capacity of mammalian PNS neurons, and silence of such genetic programs is responsible for the diminished intrinsic axon regeneration ability of mammalian CNS neurons. Our study provided clear and strong evidence that X chromosome encoded histone deacetylase Kdm6a and its paralog Kdm6b, acting as regeneration repressors to regulate spontaneous sensory axon regeneration *in vivo*. More importantly, conditional Kdm6a deletion in RGCs resulted in robust optic nerve regeneration and increased RGC survival via reshaping RGC chromatin and transcriptomic landscape back to a developmental-like state. Moreover, Klf4 was identified as a downstream target of the Kdm6a-H3K27me3 signaling and overexpression of Klf4 resulted in impaired axon regeneration. Furthermore, several additional genes known to suppress axon growth have also been identified as potential targets regulated by Kdm6a. These results suggest that in addition to many regeneration enhancing genes, the expression of axon growth repressors also play critical roles in silencing the intrinsic axon regeneration ability of mature mammalian neurons. Because H3K27me3 functions to repress gene expression, it might be a key epigenetic factor for suppressing the expression of regeneration repressor genes in mature neurons. Collectively, our study offers a novel strategy for identifying novel repressors of axon regeneration, expanding the pool of genes that can be manipulated to promote CNS axon regeneration in future studies.

To our surprise, knocking out *Kdm6b* in RGCs had little effect on optic nerve regeneration after ONC. Despite up to 84% of sequence similarity in the JmjC domain between Kdm6a and Kdm6b,[34] our results suggested that Kdm6b might not be the major H3K27me3 demethylase in RGCs. Indeed, accumulating evidence showed that Kdm6a and Kdm6b in many cases played different roles in cellular reprogramming,[43, 44] neural commitment,[74] skeletal formation,[75] embryonic development,[76, 77] muscle regeneration,[78] lifespan extension,[79] plasma cell formation,[80] and tumorigenesis.[81, 82] The distinct functional roles of Kdm6a and Kdm6b might be associated with their demethylase-dependent or - independent activities, relative expression levels, different post-translational modifications or complex formation. In our study, knocking out *Kdm6a* or knocking in *Kdm6a* mutant without demethylase activity enhanced optic nerve regeneration to the similar extent, indicating that the demethylase activity of Kdm6a was responsible for repressing axon regeneration. Based on single-cell RNA-seq results from recent study,[57] the expression level of Kdm6b is actually higher than that of Kdm6a in RGCs. Interestingly, our immunostaining results showed that Kdm6b was mainly located in the cytoplasm of RGCs, which might explain its lack of histone demethylase activity in RGCs. Indeed, evidence from other studies has shown that in addition to regulating histone methylation in the nucleus, Kdm6b might also control the methylation of non-histone proteins in both the nucleus and the cytoplasm. Moreover, the demethylase independent functions of Kdm6b have also been discovered. For instance, Kdm6b reduces the efficiency and kinetics of somatic cell reprogramming via both upregulating *INK4a/Arf* expression in a demethylase activity-dependent manner or targeting PHF20 for ubiquitination and degradation through recruiting the E3 ubiquitin ligase Trim26 in a demethylase activity- independent manner, with the latter having a predominant role.[83] Furthermore, Kdm6b has been shown to interact directly with p53, controlling its trafficking and subcellular distribution. This functional interaction regulates the retention or translocation of p53 to the nucleus, thereby modulating a number of biological processes, including neural stem cell differentiation,[84] gliomagenesis[85] and neuronal apoptosis.[86] Therefore, in our study it is very likely that Kdm6b located in the cytoplasm regulated RGC survival via an H3K27 demethylation independent manner.

Unlike that of axon regeneration, deletion of either Kdm6a or Kdm6b markedly protected RGCs from apoptosis, but via different underlying mechanisms. Thus, our study demonstrated that Kdm6a and Kdm6b played different roles with different mechanisms in regulation of optic nerve regeneration and neuroprotection, respectively (**Figure 8**). It is notable that multiple studies have demonstrated that Kdm6a and Kdm6b have both shared and distinct target genes.[82, 87, 88] The precise mechanism by which this differential target recognition is determined remains unknown. Nevertheless, it is conceivable that the involvement of their N-terminal region in distinct complexes or subcellular localizations contributes to this distinctive specificity. A recent study showed that Kdm6a downregulation conferred neuroprotective effects via the H3K27me3-mediated NOTCH2/Abcb1a axis *in vitro* and *in vivo*.[89] Similar studies of Kdm6b showed that its depletion also prevented neuronal apoptosis by a dual mechanism, with distinct downstream genes. The inhibition of Kdm6b has been demonstrated to significantly alleviate neuronal apoptosis by downregulating matrix metalloprotease (MMP) in cooperation with NF-κB,[90] as well as Bax/Caspase-3 in a p53-dependent manner.[86] Strikingly, our study further emphasized that Kdm6a/b exerted different roles by a distribution-dependent mechanism, despite the high degree of sequence similarity in their catalytic domains. Future studies are necessary to dissect out the molecular mechanisms underlying their different roles in controlling CNS neuroprotection and axon regeneration.

**Fig. 8.**
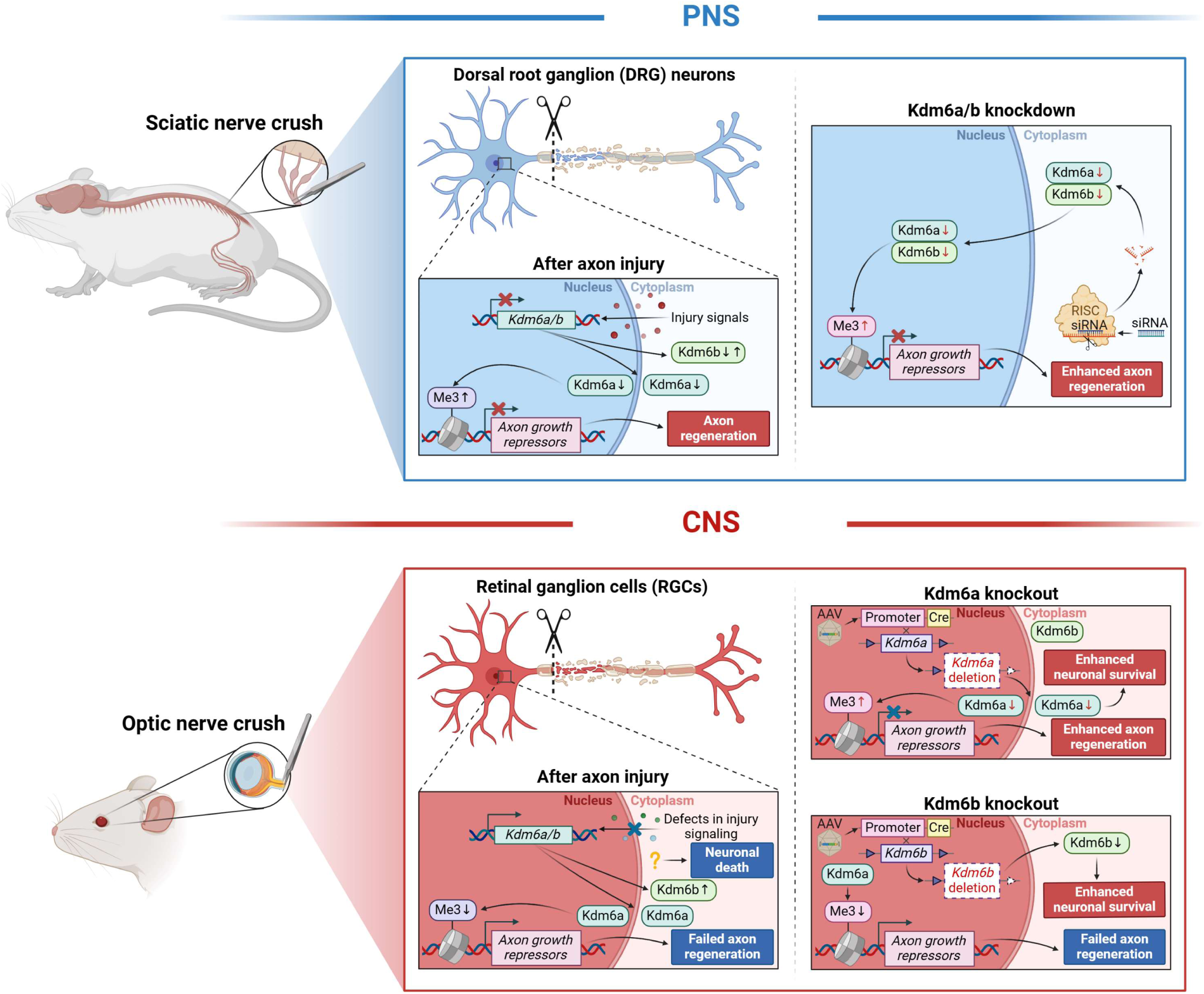
Kdm6a and Kdm6b play different roles in regulation of PNS and CNS neural regeneration due to their distinct cellular localizations. In the peripheral nervous system (PNS), sciatic nerve crush-induced injury signal leads to spontaneous axon regeneration by reshaping the transcriptomic landscape of dorsal root ganglion (DRG) neurons. During such process, the protein level of histone demethylase Kdm6a is reduced in both the cytoplasm and the nucleus, whereas the level of Kdm6b decreases specifically in the nucleus. As a result, the level of histone 3 lysine 27 trimethylation (H3K27me3) is elevated, leading to suppressed expression of axon growth-associated repressors (e.g. Klf4) and the subsequent axon regeneration. Knocking down either Kdm6a or Kdm6b before nerve injury mimics the injury signal, leading to elevated level of H3K27me3 and enhanced axon regeneration. In the central nervous system (CNS), after the optic nerve axonal injury the level of Kdm6a in either the cytoplasm or the nucleus of retinal ganglion cells (RGCs) remains the same. In contrast, Kdm6b is mainly located in the cytoplasm and its level slightly elevated after the injury. As a result, the level of H3K27me3 in RGCs is low, allowing the expression of axon growth repressors (e.g. Klf4) and axon regeneration failure. In the meantime, optic nerve injury leads to significant neuronal cell death with elusive underlying mechanism. In RGCs, only Kdm6a is localized in the nucleus. Therefore, knocking out Kdm6a is sufficient to elevate the level of H3K27me3 and enhance axon regeneration. In addition, decreased level of Kdm6a also shows neuroprotective function with enhanced neuronal survival. In contrast, knocking out Kdm6b mainly located in the RGC cytoplasm has no effect on the H3K27me3 level in the nucleus, allowing the expression of axon growth repressors and failed axon regeneration. However, downregulation of Kdm6b in the cytoplasm can enhance neuronal survival independent of its histone demethylase activity. Schematic was created with BioRender (www.biorender.com)

Although many genes have been identified to induce optic nerve regeneration, we have limited knowledge about how neuronal transcriptomics landscape is reshaped during regeneration via epigenetic modification. By performing and analyzing RNA-seq of enriched RGCs at different developmental/maturation stages and between wild type and *Kdm6a* knockout mice, we found that deleting *Kdm6a* in mature RGCs changed their transcriptomic landscape back to a developmental-like state. As a result, mature RGCs regained their intrinsic ability to support axon regeneration. In our latest study,[30] we found that H3K27 methyltransferase Ezh2 in post-mitotic neurons functioned to repress dendrite development during development. Together with this study, it seems that H3K27me3 functions as a major epigenetic regulator of neuronal morphogenesis, promoting axon growth but suppressing dendrite development. Importantly, epigenetic regulation, such as histone modifications, can orchestrate a large-scale change in gene expression to regulate biological functions. The Kdm6a signaling revealed in this study are well positioned to reshape the gene expression landscape of mature CNS neurons at both the transcriptional and translational levels to enhance axon regeneration.

In our study, we also provided evidence that Kdm6a regulated optic nerve regeneration via distinct mechanism from that of Pten, independent of mTor activation. Moreover, combining Kdm6a and Pten double deletion resulted in long-distance axon regeneration passing the optic chiasm. 3D imaging of tissue-cleared optic nerve showed that regenerating axons induced by double deletion have more streamlined trajectory and increased branching, indicating enhanced growth ability. At the border of the optic chiasm, more than half axons either stopped or turned back, highlighting the inhibitory nature of the chiasm microenvironment. For axons entering the chiasm, many of them projected into the ipsilateral optic tract, indicating the lack of proper axon guidance. Therefore, for long-distance regenerating axons to re-innervating their original targets for functional recovery, modifying the microenvironment of the chiasm and/or the guidance sensing receptors in the regenerating axons would be major near-future research focus in the field.

Lastly, Kdm6a has been shown to escape from X inactivation and be expressed in the inactivated X chromosome. As a result, the protein level of Kdm6a is higher in female than that in male mice in most cells, including neurons. A recent study[91] showed that Kdm6a contributed to the resistance of neurodegenerative insults of neurons in female mice. Interestingly, based on previous epidemiology studies, it is known that women are more likely to be affected by glaucomatous disease, especially for angle-closure glaucoma (ACG), for which nearly 70% are women.[92, 93] Our study raised the possibility that a higher level of Kdm6a in female RGCs might increase the susceptibility of female animals to retinal injury.

## 4. Experimental Section

### 4.1. Animals

All experiments involving animals were performed in accordance with the animal protocol approved by the Institutional Animal Care and Use Committee of the Johns Hopkins University, Chinese Academy of Sciences and Zhejiang University. 8- to 12-week old adult female CF-1 mice (weighing from 30 to 35 g) were purchased from Charles River Laboratories and housed in the University Animal Facility. The female *Kdm6a^f/f^* mutant mice (Kdm6atm1c(EUCOMM)Jae) possessing loxP sites flanking exon 3 of the Kdm6a (lysine (K)- specific demethylase 6A) gene, the *Kdm6b^f/f^* mutant mice (B6.Cg-Kdm6btm1.1Rbo/J) possessing loxP sites flanking exons 14-20 of the Kdm6b (lysine (K)-specific demethylase 6B) gene and the *Pten^f/f^* mutant mice (B6.129S4-Ptentm1Hwu/J) possessing loxP sites flanking exon 5 of the Pten (phosphatase and tensin homolog) gene were purchased from The Jackson Laboratory. *Kdm6a* enzyme-dead knockin mice possessing the H1146A and E1148A point mutations in exon 24, as a gift, were obtained from Dr. Kai Ge at the National Institutes of Health.

### 4.2. Immunostaining, fluorescence microscopy, and image analysis

The DRG or retina cryostat sections were collected and immunostained with the antibody against H3K27me3 (1:1000), Kdm6a (1:500), Kdm6b (1:500), or p-S6 (1:500) each co-immunostained with the neuronal marker Tuj1 (1:1000) antibody. To quantify the fluorescence intensity of H3K27me3, Kdm6a, Kdm6b, or p-S6 in sensory neurons, only Tuj1 positive cells were selected for measurement. The mean fluorescence intensity of each sensory neuron in DRG or RGC in retina section was measured using the ImageJ software. The percentage of p-S6 positive RGCs was calculated by dividing the number of p-S6/Tuj1 double positive cells by the number of Tuj1 positive cells. For DRG, 3 independent DRG tissues in each condition were used in each condition. For each DRG, 3 sections were selected and 30 sensory neurons in each section were selected for measurement. For retina, 3 independent retinal tissues in each condition were used. For each retina, 3-8 sections were selected and 15-20 RGCs in each section were selected for measurement. The “n” in the figures indicates the number of independent experiments.

### 4.3. In vivo electroporation of adult DRG neurons and quantification of axon regeneration

The in vivo electroporation of adult mouse DRGs was performed as described previously.[56] Briefly, under anesthesia induced by ketamine (100 mg/kg) and xylazine (10 mg/kg), a small dorsolateral laminectomy was performed to expose the L4 and L5 DRGs. EGFP plasmid or EGFP plus siRNA oligos or expression plasmid (1-1.5 μl per ganglion) were injected into the DRGs using pulled glass capillaries and a Picospritzer II (Parker Ins.; pressure, 30 psi; duration: 8 ms). Immediately after injection, electroporation was performed by applying five pulses of current (35 V, 10 msec, 950-msec interval) using a custom-made tweezer-like electrode powered by the Electro Square Porator ECM830 (BTX Genetronics). The mice were allowed to recover after closing the wound. Two days after the electroporation, the sciatic nerves were crushed with fine forceps and the crushed sites were marked with nylon epineural sutures (size 10-0). Two days later, the mice were perfused with ice-cold 4% PFA in sodium phosphate buffer (pH 7.4). The whole nerve segment was then dissected out and further fixed in 4% PFA overnight at 4°C. Before whole-mount flattening, it was confirmed that the place of epineural suture matched the injury site, and experiments were included in the analysis only when the crush site was clearly identifiable.

For quantification of in vivo axon regeneration, the fluorescence images of the whole mount nerves were first obtained. All identifiable EGFP-labeled axons in the sciatic nerve were then manually traced from the crush site to the distal growth cone to measure the length of axon regeneration. Only nerves with at least 15 identifiable axons were measured, and the “n” in the figures indicates the number of mice.

### 4.4. Quantitative real time polymerase chain reaction (Real-time PCR)

Total RNA was isolated with the TRizol Reagent (Thermo Fisher Scientific), then reverse transcribed by using the M-MLV reverse transcriptase (Roche Applied Science). To quantify the mRNA levels with real-time PCR, aliquots of single-stranded cDNA were amplified with gene-specific primers and Power SYBR Green PCR Master Mix (Invitrogen) using the CFX96™ real-time PCR detection system (Bio-Rad). Specific primers used in this study are listed in **Extended Data table 1**. The PCR reactions contained 20-40 ng of cDNA, Universal Master Mix (Invitrogen), and 200 nM of forward and reverse primers in a final reaction volume of 20 μl. The ratio of different samples was calculated by the build in data analysis software of the CFX96™ real-time PCR system.

### 4.5. Intravitreal injection, optic nerve crush, and RGC axon anterograde labeling

The mice were anaesthetized with intra-peritoneal injection of a mixture of ketamine (100 mg/kg) and xylazine (10 mg/kg). 1 μl of AAV2 viruses (Titers > 1x10^13^ vg/ml), GSK-J4 (300 mM), 1.5 μl NMDA (20 mM) for NMDA-induced retinal neurotoxicity were injected into the vitreous body of 6- to 8-week old mutant or wild type mice by glass micropipette connected to a Picospritzer II (Parker Inc.) (pressure: 15 psi; duration: 6 ms). Two weeks after intravitreal injection, the right optic nerve was exposed intraorbitally and crushed with Dumont #5 forceps for 2 seconds approximately 1 mm behind the optic disc.

To label RGC axons in the optic nerve by anterograde labeling, 1.5 μl of cholera toxin β subunit (CTB) conjugated with fluorescence Alexa-594 (2 μg/μl, Invitrogen) was injected into the vitreous 2 days before sacrificing the animals. Sacrificed animals were fixed by perfusion with 20 mL of 0.1 M phosphate buffered saline (PBS), followed with 40 mL of 4% paraformaldehyde (PFA) at 5 ml/minute. Retina and optic nerve segments were dissected out and post-fixed in 4% PFA overnight at 4 °C. The fixed optic nerves were rinsed with cold PBS three times and stored in cold PBS until further use.

### 4.6. Tissue dehydration and clearing

Optic nerves were first dehydrated in increasing concentrations of tetrahydrofuran (THF, 50%, 70%, 80%, 100% and 100% for 20 minutes each) in amber glass bottles as described in a previous publication.[94] Incubations were carried out on an orbital shaker at room temperature. Optic nerves were then transferred into benzyl alcohol/benzyl benzoate (BA:BB, 1:2 in volume; Sigma) clearing solution for 20 minutes. During the whole procedure optic nerves were protected from light to reduce the loss of CTB fluorescent signal.

### 4.7. Imaging and quantification of RGC axon regeneration in the optic nerve

The whole-mount cleared optic nerves were imaged using a Zeiss LSM 710 confocal microscope controlled by ZEN software. A 20X objective was used to acquire image stacks with 2-μm z spacing. The Motor-driven XY scanning stage with tiling function were used to scan the optic nerve with 15% overlap in the X Y dimensions.

For quantification of optic nerve axon regeneration, every 4 consecutive images were projected to generate a series of Z-projection images of 8-μm optical sections. In each Z-projected image, the number of CTB-labeled fibers was counted at 250-μm intervals distal to the crush site till the place where no fluorescence signals were visible. The numbers at each 250- μm intervals were summed over all Z-projection images.

For quantification of the average lengths of top 5 longest regenerating axons, all acquired images were merged together using ZEN software to generate the maximum intensity Z-projection images. Then the lengths of the top 5 longest axons were manually measured from the ends to the crush site with ImageJ software (NIH).

### 4.8. Analyses of axonal tip morphology and axon trajectory

The analysis of axonal tip morphology was performed as described previously.[95] For quantification of the size of the distal axon ends, the tips of top 20 longest regenerating axons were identified in the Z-projection images of each nerve. Using ImageJ software (NIH), the maximal width of the axonal tips was measured and was presented as a ratio with respect to the diameter of the adjacent axon shaft (tip/axon shaft ratio). The axon ends with tip/axon shaft ratio > 4 were defined as retraction bulb, while the others as growth cone.

For quantification of the U-turn rate, in the Z-projection images the trajectories of top 20 longest axons within optic nerve or all axons entering the optic chiasm were tracked near the axonal ends. The axonal trajectory that made a turning with the angle between the direction of the axonal tip and the longitudinal axis of the optic nerve greater than 90 degrees was defined as a U-turn. The U-turn rate of each nerve was presented by U-turn number/axon number.

For quantification of branching index, in the Z-projection images the axonal tips number was counted from the selected site (with 20 regenerating axons passing or the optic nerve-chiasm transition zone) till the place where no tips were visible. The branching index of each nerve was calculated by axonal tips number/selected axon number (20 axons for optic nerve or all axons for the optic chiasm).

### 4.9. Quantification of RGC transduction and survival rates

Retinas were dissected out for whole-mount preparations. Retinal whole-mounts were blocked in the staining buffer containing 2% BSA and 0.5% Triton X-100 in PBS for 1 hour before incubation with primary antibodies. Primary antibodies used: Tuj1 (1:500), Cre (1:500).

Retinas were incubated with primary antibodies overnight at 4°C and washed four times for 10 minutes each in PBS containing 0.5% Triton X-100. For secondary antibodies, Alexa Fluor- 488/647-conjugated antibodies were used. Confocal images were acquired using a Zeiss LSM 710 confocal microscope (20X objective). Images were organized and analyzed using ZEN and ImageJ software. For RGC survival quantification, whole-mount retinal tissues were immunostained with Tuj1 antibody to label the survival RGCs. 15-20 fields were randomly sampled from the peripheral regions of each retina. The survival rate was calculated by measuring the ratio of Tuj1 positive cell number in retina with optic nerve crush (ONC) to that in control retina without ONC. To quantify RGC transduction efficiency by AAV2-Cre, whole-mount retinal tissues were double-immunostained with Tuj1 and Cre antibodies. 8-12 fields were randomly sampled from the peripheral regions of each retina. The transduction efficiency was calculated by counting and calculating the ratio of Cre/Tuj1 double-positive cell number to the number of Tuj1 positive cells.

### 4.10. Western blot

Total protein was extracted with the RIPA buffer containing protease inhibitor cocktail and phosphatase inhibitor cocktail. 15 μg of each protein sample was separated by a 4-12% gradient SDS-PAGE gel and electrotransferred onto polyvinylidene fluoride membranes. Nonspecific bands were blocked with 5% non-fat milk in 1x TBS containing 0.1% tween-20 (TBST) for 1 hour at room temperature. Membranes were incubated with the primary antibodies of Kdm6a (1:1000), Kdm6b (1:1000), β-actin (1:10000) as internal control overnight at 4 °C and followed by HRP-conjugated secondary antibodies with a concentration of 1:2000 for 1 hour at room temperature. Membranes were washed with TBST for 15 minutes three times after each antibody incubation. The immunocomplex bands were visualized by the ECL kit (GE Healthcare).

### 4.11. Purification of RGCs

Retinas were dissected, incubated with 20 units/ml papain and 0.005% DNase (Worthington) for 10 minutes at 37 °C, then dissociated into a single-cell suspension in pre-sort buffer (BD FACS). Retinal cells were filtered through a 40-μm cell strainer (BD Falcon), then collected by centrifugation at 500 g for 5 minutes. Supernatant was removed and immediately pelleted cells were resuspended in 400 μl of pre-sort buffer. Cell suspension was incubated with Fc block antibodies for 5 minutes at 4°C and then stained with Thy1.2-FITC antibody and IgG2a isotype antibody (Thermo fisher) as a negative control for 30 minutes at 4 °C, respectively. Retinal cells were pelleted by centrifugation at 500 g for 5 minutes. Supernatant was discarded and pelleted cells were washed one time with 500 μl of pre-sort buffer. Cell suspension was incubated with DAPI for 5 minutes before FACS sorting, and then sorted on a BD FACSAria Fusion using a 100-micron nozzle. Sorted cells were immediately subjected to subsequent experiments.

For RGC enrichment by immunomagnetic cell separation (MACS), cell suspension was incubated with Fc block antibodies for 5 minutes at 4 °C and then labeled with anti-mouse Thy1.2 magnetic particles (BD IMag™) for 30 minutes at 4 °C. Tubes were placed on the cell separation magnet at room temperature for 8 minutes (BD IMag™), then the supernatant was aspirated off carefully for leaving unlabeled cells. Labeled cells were resuspended in 1x BD IMag™ buffer and the tubes were returned to the magnet for another 4 minutes. The separation was repeated twice to increase the purity of the positive fraction. After the final wash step, purified cells were immediately used for further analyses.

### 4.12. Library preparation and RNA sequencing

A total amount of 2 μg RNA per sample was used as input material for the RNA sample preparations. Sequencing libraries were generated using NEBNext® Ultra™ RNA Library Prep Kit for Illumina® (#E7530L, NEB, USA) following the manufacturer’s recommendations and index codes were added to attribute sequences to each sample. Briefly, mRNA was purified from total RNA using poly-T oligo-attached magnetic beads. Fragmentation was carried out using divalent cations under elevated temperature in NEBNext First Strand Synthesis Reaction Buffer (5X). First strand cDNA was synthesized using random hexamer primer and RNase H. Second strand cDNA synthesis was subsequently performed using buffer, dNTPs, DNA polymerase I and RNase H. The library fragments were purified with QiaQuick PCR kits and elution with EB buffer, then terminal repair, A-tailing and adapter added were implemented. The aimed products were retrieved and PCR was performed, then the library was completed.

RNA concentration of library was measured using Qubit® RNA Assay Kit in Qubit® 3.0 to preliminary quantify and then dilute to 1 ng/μl. Insert size was assessed using the Agilent Bioanalyzer 2100 system (Agilent Technologies, CA, USA), and qualified insert size was accurate quantification using StepOne Plus™ real-time PCR System (Library valid concentration > 10 nM). The clustering of the index-coded samples was performed on a cBot cluster generation system using HiSeq PE Cluster Kit v4-cBot-HS (Illumina) according to the manufacturer’s instructions. After cluster generation, the libraries were sequenced on an Illumina platform and 150 bp paired-end reads were generated.

### 4.13. Bulk RNA-seq data analysis

Using bulk RNA-seq, we measured gene expression profiles of AAV-Cre (Kdm6a), AAV-GFP (wild type), and postnatal day (P) 1, P14, P21 samples, with at least two biological replicates for each condition or timepoint. The bulk RNA-seq data were obtained from Gene Expression Omnibus (GEO: GSE244243). All raw sequencing reads were quality checked with FastQC (v0.11.9), and leading bases were trimmed from the reads with Trimmomatic (v0.39) as appropriate. Quality-checked reads were mapped to the GRCm38/mm10 mouse reference genome with splice-aware aligner HISAT2 (v2.2.1). FeatureCounts (Subread v2.0.3) was used to estimate the expression counts of the genes.

For differential gene expression analysis, we used DESeq2 (v1.34.0) to perform normalization and differential expression test on the raw read counts for each gene. Differential expression genes (DEGs) were defined using DESeq2 with the adjusted *P*-value < 0.05 and |log2(Fold Change)| > 1. The gene ontology (GO) enrichment analysis was performed by clusterProfiler (v4.2.2). In order to integrate and analyze Kdm6a knockout data and optic nerve development data, the TPM (transcriptsper million reads) matrix for genes and transcripts calculated by RSEM (v1.3.3) were used. We also used the ComBat in sva (v3.42.0) to minimize the technical effect of different data. The corrected values were subjected to principal components analysis (PCA) and hierarchical clustering.

### 4.14. Single-cell RNA-seq data analysis

The data used to analyze the time course of transcriptomic changes in single retinal ganglion cells following ONC was downloaded from Gene Expression Omnibus (GEO: GSE137398).[57] Then, the expression matrices were integrated by Seurat (v4.1.0) with default parameters. The changes of average expression levels of *Kdm6a* and *Kdm6b* in all RGCs were mainly analyzed at different time point after ONC, as showed in dot plot and violin plot. The data used to analyze the expression levels of all Klf family members (*Klf1-18*) in adult sensory neurons after SNI was downloaded from Gene Expression Omnibus (GEO: GSE154659).[73] Dot plot showed the changes of average expression levels of all Klf family members in all sensory neurons at day 3 after SNI.

### 4.15. CUT&Tag

Approximately 1x10^5^ RGCs purified by coupling Thy 1.2 beads were processed by centrifugation, and were washed with Wash Buffer (20 mM HEPES, pH 7.5; 150 mM NaCl; 0.5 mM Spermidine; 1X Protease inhibitor cocktail; 0.05% DMSO-Digitonin). The supernatant was removed and the bead-bound cells were resuspended with 50 µL Dig-wash Buffer (20 mM HEPES, pH 7.5; 150 mM NaCl; 0.5 mM Spermidine; 1x Protease inhibitor cocktail; 0.05% DMSO-Digitonin) containing 2 mM EDTA and 1% BSA and a 1 : 50 dilution of the primary antibody. Primary antibody incubation was performed on a rotating platform for 2h at room temperature. The primary antibody was removed by transient centrifugation and placing the tube on the magnet stand to clear and pulling off all the liquid. The appropriate second antibody was diluted 1 : 50 in 50 µL Dig Wash Buffer and bead-bound cell were incubated for 1h at room temperature. Cells were washed using the magnet stand 2 times for 5min in 200 µL Dig-Wash buffer to remove unbound antibodies. A 1 : 200 dilution of pG-Tn5 adapter complex (∼0.04 µM) was prepared in Dig-300 Buffer (0.05% DMSO-Digitonin, 20 mM HEPES, pH 7.5, 300 mM NaCl, 0.5 mM Spermidine, 1x Protease inhibitor cocktail). After removing the liquid on the magnet stand, 100 µL was added to the cells with gentle vortexing, which was incubated with pG-Tn5 at RT for 1 h. Cells were washed 2 times for 5min in 200 µL Dig-300 Buffer to remove unbound pG-Tn5 protein. Next, cells were resuspended in 300 µL Tagmentation Buffer (10 mM MgCl 2 in Dig-300 Buffer) and incubated at 37 °C for 1h. To stop tagmentation, 2.25 µL of 0.5 M EDTA, 2.75 µL of 10% SDS and 0.5 µL of 20 mg/mL Proteinase K was added to 300 µL of sample, which was incubated overnight at 37 °C. To extract the DNA, 300 µL PCI (Phenol : Chloroform : Isoamylol = 25 : 24 : 1) were added to each sample with vortexing, quickly spun 5s ,16000g for 15 min at room temperature. Absorbed the upper water phase to a new 1.5 mL tube and added 300 chloroform, 16000 g for 15 min at room temperature. Absorbed the upper water phase to a new 1.5 mL tube and added 700 µL 100% ethyl alcohol, 16000g for 15 min at 4 °C. Pulled off all the liquid and added 1 mL 100% ethyl alcohol 16000g for 1 min at 4 °C.

Carefully discarded all the liquid and aired the tube 5 min, 25 µL 1X TE, 10 mM Tris-HCl, pH 8.0) containing 1 mM EDTA were added to dissolve the DNA. To amplify libraries, 24 µL DNA was mixed with 5 µL a uniquely index i7 and i5 primer; Using a different index for each sample. A volume of 16 µL PCR Master mix was added and mixed. The sample was placed in a Thermocycler with a heated lid using the following cycling conditions: 72 °C for 3 min ; 98 °C for 30 s; 17 cycles of 98 °C for 10 s and 60 °C for 30s and 72 °C for 30s; final extension at 72 °C for 5 min and hold at 4 °C. Post-PCR clean-up was performed by adding 1.1× volume of DNA clean beads (Nanjing, vazyme), and libraries were incubated with beads for 15 min at RT, washed twice gently in 80% ethanol, and eluted in 20 µL sterile water.

### 4.16. Chromatin Immunoprecipitation (ChIP) assay

ChIP assay of DRG tissue was conducted in accordance with the protocol provided by the Active Motif company, with a few modifications. Briefly, 6-8 naïve or 10-15 axotomized L4 and L5 DRGs were collected and homogenized with 1% formaldehyde (Sigma-Aldrich) for 20 min. The homogenized tissue were washed with cold PBS, suspended with 500 μl cold cell lysis buffer (5mM PIPES, pH 8.0, 85mM KCl, 0.5% NP40, and 1X complete proteinase inhibitor), and then incubated on ice for 5 min. The lysates were centrifuged at 3000rpm for 5 minutes, and the pellets were resuspended in 800μl of SDS Lysis Buffer (Millipore). After 20- minute incubation on ice, lysates were sonicated (6 pulses, 10 seconds each at a power output of 40%, with 1-minute incubations on ice in between each pulse) to shear the genomic DNA into 200 and 1000 base pair fragments. To verify the size of the sheared chromatin (average size ∼500-600bp), 5 μl aliquots of the lysates were treated with 1μl of Proteinase A (20 mg/ml) for 20 min at 50°C and the sample was analyzed using a 1.5% agarose gel.

To perform the immuno-precipitation, the sonicated cell supernatant was mixed with protein G Magnetic Beads, ChIP buffer 1, protease inhibitor cocktail, water and 5μg of normal rabbit IgG or rabbit anti-H3K27me3 antibody in 1.7ml tubes. The chromatin was rotated overnight at 4°C. Next day the beads were washed one time with 800 μl ChIP Buffer 1 and two times with 800 μl ChIP buffer 2. After the final wash, the supernatant was removed as much as possible without disturbing the beads. The washed beads were treated with the elution buffer, followed by reverse cross-linking buffer. Formaldehyde-induced protein-DNA crosslinking were heat reversed by incubating protein-DNA complex at 65°C overnight. The obtained DNA was then treated with 2 μl proteinase K for 1 hour, followed by 2 μl proteinase K stop solution. Briefly centrifugation was performed to collect the purified DNA fragments, which could be used immediately in PCR or stored at -20°C.

### 4.15. Statistics

Statistics were analyzed using GraphPad Prism 7.0 and the significance was set at a value of p < 0.05. Data are presented as mean ± SEM. For two group comparisons, if the data were normalized to the control condition and presented as percentage of control, one sample t test was used to determine the statistical significance. Otherwise, the regular two tailed student’s t test was used. For comparison between multiple groups, the one-way ANOVA followed by Tukey’s multiple comparison test was used to determine the statistical significance. The “n” indicates the number of independent experiments unless specifically stated otherwise.

## Contributions

S-G. Y., C-M. L., and F-Q. Z. initiated the projects and designed the experiments. S-G. Y., X-W. W., and C-P. L. performed most of experiments and analyzed data. T. H. performed most of the bioinformatics analyses. C. Q., Q. L., L-R. Z., S-Y. Z., C-Y. D., and S. performed experiments and analyzed the data. S-G. Y. and F-Q. Z. wrote the manuscript with inputs from all the authors.

## Acknowledgements

The study was supported by grants (to F-Q.Z.) from NIH (R01NS085176, R01EY027347, R01EY030883, R01EY031779), the Craig H. Neilsen Foundation (259450), the BrightFocus Foundation (G2017037), the Pioneer and Leading Goose R&D Program of Zhejiang Province, China (2024C03028), and the Leading Innovation and Entrepreneurship Team Program of Zhejiang Province, China (2023R01005). C-M.L. was supported by the National Science Foundation of China (No. 81571212).

